# Microbial community structure in rice, crops, and pastures rotation systems with different intensification levels in the temperate region of Uruguay

**DOI:** 10.1101/2021.05.24.445164

**Authors:** Sebastián Martínez

## Abstract

Rice is an important crop in Uruguay associated mostly with livestock production in a rice and pasture rotation system since the 1920s. However, in recent years there has been interest in intensifying the production in some of these systems to satisfy market demands and increase income. Intensification occurs by augmenting the rice frequency in the rotation, including new crops like sorghum and soybean, or shortening the pasture phase. A long-term experiment was established in 2012 in the main rice producing area of Uruguay with the objective to study the impact of intensification in rice rotations. After the first cycle of rotation soils from seven rotation phases were sampled and microbial communities were studied by means of high-throughput sequencing of Illumina NovaSeq 6000. Archaeal/bacterial and fungal community composition were studied (16S rRNA and 18S gene regions) detecting 3662 and 807 bacterial and fungal Operational Taxonomic Units (OTUs), respectively. *Actinobacteria*, *Firmicutes* and *Proteobacteria* were the most common bacterial phyla. Among them, only *Proteobacteria* differed significantly between rotations. Although most fungal OTUs were unidentified, *Ascomycota*, *Basidiomycota* and *Mucoromycota* were the most abundant fungal classes within identified taxa. Bacterial communities differed between rotations forming three groups according to the percentage of rice in the system. Fungal communities clustered in four groups, although not well differentiated, and mostly associated with the antecessor crop. Only P and C:N varied between rotations among soil physicochemical variables after six years, and individual bacterial OTUs appeared weakly influenced by P, pH, Mg and fungal OTUs by P. The results suggest that after six years, bacteria/archaeal communities were influenced by the time with rice in the rotation, and fungal communities were more influenced by the antecessor crop. More studies are needed to associate fungal communities with certain rotational or environmental variables. Some taxa were associated with a particular rotation, and some bacterial taxa were identified as biomarkers. Fungal indicator taxa were not identified at the species level for any rotation.

## 1. Introduction

Soil microorganisms mediate different and important ecosystem functions associated with nutrient cycling, primary productivity, contaminant remediation, C cycle processes and climate regulation (Bardgett and van der Putten, 2014). Soil microbial diversity and processes are critical for maintaining sustainability and production of diverse agricultural systems (Bending et al. 2004; Bünemann et al. 2018). In addition, soils are reported to substantially impact microbial community diversity and structure (Bending et al. 2004; McGuire and Treseder 2010). Several soil physicochemical and biological factors are reported to influence the structure and diversity of soil microbiomes, including pH, water availability, soil type, soil texture, weather conditions, interactions with other underground and aboveground soil organisms, as well anthropogenic, geological and historical factors (Bending et al. 2004; Brockett et al. 2012; Zheng et al. 2019). Agriculture is reported to cause disturbances in soils, particularly affecting some of the physicochemical factors mentioned previously and, therefore, the associated microbiota (Bending et al. 2004).

Rice (*Oryza sativa* L.) is one of the most cultivated crops feeding more than 50% of the world population and influencing the livelihoods and economies of several billion people (IRRI, 2006). Soils subjected to irrigated rice cultivation differ from most other crop soils in that they are subjected to oxic and anoxic phases within the crop cycle. These changes result in the selection of specific physiological groups of microorganisms correlated with chemical differences owing to their metabolism, being, aerobic, anaerobic, or facultative (Brune et al., 2000). The physicochemical properties show great variation among soil types found in different geographical locations, and, thus, soil microbial communities change accordingly. Most reports focus on the changes among microbial communities in soils with different nutritional statuses (Jiang et al., 2016).

Several studies on microbial communities in paddy soils focused on the effects of crop rotation, fertilization and residue retention on the soil microbial abundance, biomass, composition and activity (Kim and Lee, 2020). But, most of these studies focused on rice production systems in Asia where continuous rice, with two or three crops each year, is substituted by rotations of rice with other crops, such as wheat, pulse, oilseeds, maize and vegetables (Hoa et al., 2006; Goswamia et al., 2020). However, the structures and patterns of microbial communities in paddy soils are poorly characterized across systems, soils and crop rotation gradients in non-Asiatic countries and/or temperate regions.

Changes in community composition, diversity and networks are most determined by the crop rotation when comparing rice monoculture with rice systems including legumes and other crops. Paddy soils in temperate to tropical regions of Asia support bacterial communities mostly driven by soil characteristics and fungal communities are better predicted by geographical distance (Jiang et al. 2016). Abundance, composition, and diversity of bacterial communities are very different in rotations of rice with other crops, like mungbean, wheat, or maize than rice in monoculture (Jiang et al. 2016; Feng et al. 2020). Rice fertilization type also produces differential effects on microbial communities, decreasing fungal richness and increasing bacterial richness when organic manure is added or increasing microbial community abundance under rice-wheat rotation compared with rice fallow (Chen et al. 2016; Feng et al. 2020). Moreover, bacterial diversity is decreased in soils receiving straw incorporation with high fertilization inputs, which denotes that higher C:N ratio is better for bacterial diversity and depends on the affected bacterial phyla (Wu et al. 2020). Also, in rice-wheat crop rotation with different fertilization inputs, microbial communities respond differently. Bacterial community composition and functions were modified by fertilization only, while fungal community composition was affected by both fertilization and crop types (Chen et al. 2020). However, little is known about how different rice rotations affect soil microbial communities and how it affects the productivity and sustainability of these systems in temperate regions like Uruguay. Previous studies showed no major differences among microbial communities associated with regards to rice and pasture rotations and soils under continuous rice in South America (Fernández et al., 2013; Maguire et al., 2020).

Diversity of microbial communities, either bacterial or fungal, inhabiting the rice soil ecosystem have been characterized previously, both in anoxic bulk soil and in oxic surface soil (Lee et al., 2015; Kim and Lee, 2019). Most studies reported bacterial phyla *Proteobacteria*, *Chloroflexi*, *Actinobacteria* and *Acidobacteria* and fungal classes *Ascomycota*, *Basidiomycota* and *Glomeromycota* dominating rice paddy soils (Jiang et al., 2016). In Uruguayan conditions, *Firmicutes*, *Proteobacteria*, and *Acidobacteria* were dominating soils under traditional rice and pasture rotation with high relative abundance of *Firmicutes* (~40%). Other phyla were found with <10% relative abundance, except for *Verrucomicrobia* and *Actinobacteria* that incremented relative abundances in the soil after four years or under permanent pasture (Fernández et al., 2013). Uruguayan native grassland soils of the Campos biome, where rice agroecosystems are established, were dominated by *Proteobacteria*, *Actinobacteria*, *Firmicutes*, *Verrucomicrobia*, *Acidobacteria*, *Planctomycetes* and *Chloroflexi* phyla (Garaycochea et al., 2020).

Uruguayan rice system has been maintained since the 1920s in a traditional rotation of rice and pastures, consisting of two years of rice and three to four years of perennial pastures in legume–-grass mixtures integrated with livestock production (Chebataroff, 2012). Cultivation is exclusively under irrigation with a dry phase from seeding to approximately one month later when it is flooded and completes the cycle until harvest (Chebataroff, 2012). Rice is grown after a perennial grass and legume mix pasture that can be beneficed by the over 100 kg N ha^-1^ fixed in a 26-week period reported for a mixture of white clover and fescue (Labandera et al., 1988). This integration of rotation systems with crops and pastures is rare globally, except in some parts of South America and particularly in Uruguay where most crop lands were historically occupied by grasslands (García-Préchac et al., 2004). This rice–pasture integration is a more sustainable system that allows preservation of natural resources, minimization of agrochemical inputs, and reduction of environmental impacts while sustaining economic productivities (Pittelkow et al., 2016). However, in recent years, there has been interest to intensify production in these systems to meet market demands and increase farmers’ incomes. Intensification occurs in different ways, like augmenting the rice frequency in the rotation with new crops and/or shortening the pasture phase. Thus, new rotations are proposed and are under continuous evaluation by farmers. Nevertheless, it is unclear whether these more intensified systems are productively and environmentally sustainable over time. To answer some of the questions that arise about intensification processes, a long-term experiment (LTE) was established in 2012 in the Experimental Unit of Paso de la Laguna (UEPL), Treinta y Tres, Uruguay. The main objectives of the experiment are to evaluate and compare the effects of intensification of rice rotations in terms of productivity, inputs use, as well as environmental indicators. Information generated during the evaluation of the experiment allows the generation of guidelines for sustainable agricultural intensification for Uruguayan rice systems (Terra et al., 2014).

In the current study, the LTE of UEPL on rice rotations was used to test the impacts of five different rotations on bulk soil microbial communities, after a complete rotation cycle of six seasons, compared with the traditional rice and pasture system. With the hypothesis that crop rotation would have an impact on the structure of the soil bacterial and fungal community, this study analyzed which antecessor of rice had the greatest impact on soil microbial communities by comparing the different phases of the experiment and determining which species can be considered indicators for these different systems. Overall, the goal of this study was to draw a baseline of the effects of crop intensification on the composition of soil microbial communities in rice fields in the temperate weather of Uruguay.

## 2. Materials and methods

### 2.1 Experimental site

The study was conducted at the LTE of the UEPL established in 2012 at Treinta y Tres, Uruguay (S 33° 16’ 22”, W 54° 10’ 23”). The field experiment was an LTE for the study of rice rotations with pastures and other crops like soybean and sorghum. The main objectives of the LTE were to study sustainability of a medium to long time of intensified rice systems in terms of productivity (Macedo et al., 2021). According to the description in Onofri et al. (2016), the LTE contain several rotations of different lengths and different numbers of phases per crop and rotation cycle and is maintained with complete straw returning and no-tillage (Macedo et al., 2021). The experiment consisted of six different rice rotations with different degrees of rice and crop intensification, from a traditional Uruguayan rice–pasture system, consisting of two consecutive seasons with rice and three years of a pasture with a mixture of grasses and leguminous species (Deambrosi et al., 2009), to a very intensified system of continuous rice (Fig. S1). Although continuous rice is a common practice in several Asiatic countries, it is unpracticed in Uruguay. The six rotations are from one to six years long with each phase occurring each year for each of the rotations, totaling 20 phases; nine of these phases are with rice each year (Fig. S1). Each phase was seeded in a 20 x 60 m plot and repeated three times (blocks) for a total of 60 plots. Rotations included: 1) continuous rice (CR, 1 year), 2) rice and other crops (RC, rice-soybean-rice-sorghum, 4 years), 3) rice and short pasture (RSP, rice-pasture, 2 years), 4) traditional rice and pasture (RP, rice-rice and three years of pasture, 5 years), 5) rice, soybean and pastures (RCP, rice-soybean-soybean-rice and pasture, 6 years), and 6) rice and soybean (RS, rice-soybean, 2 years; Fig. S1). All rotations/phases completed a full cycle in the 2017-2018 season and began a new one in the season 2018-2019 when sampling for the present study was performed before rice seeding. From the total of nine rice phases present each year, seven were selected for soil sampling (Fig. 1).

**Fig. 1.**
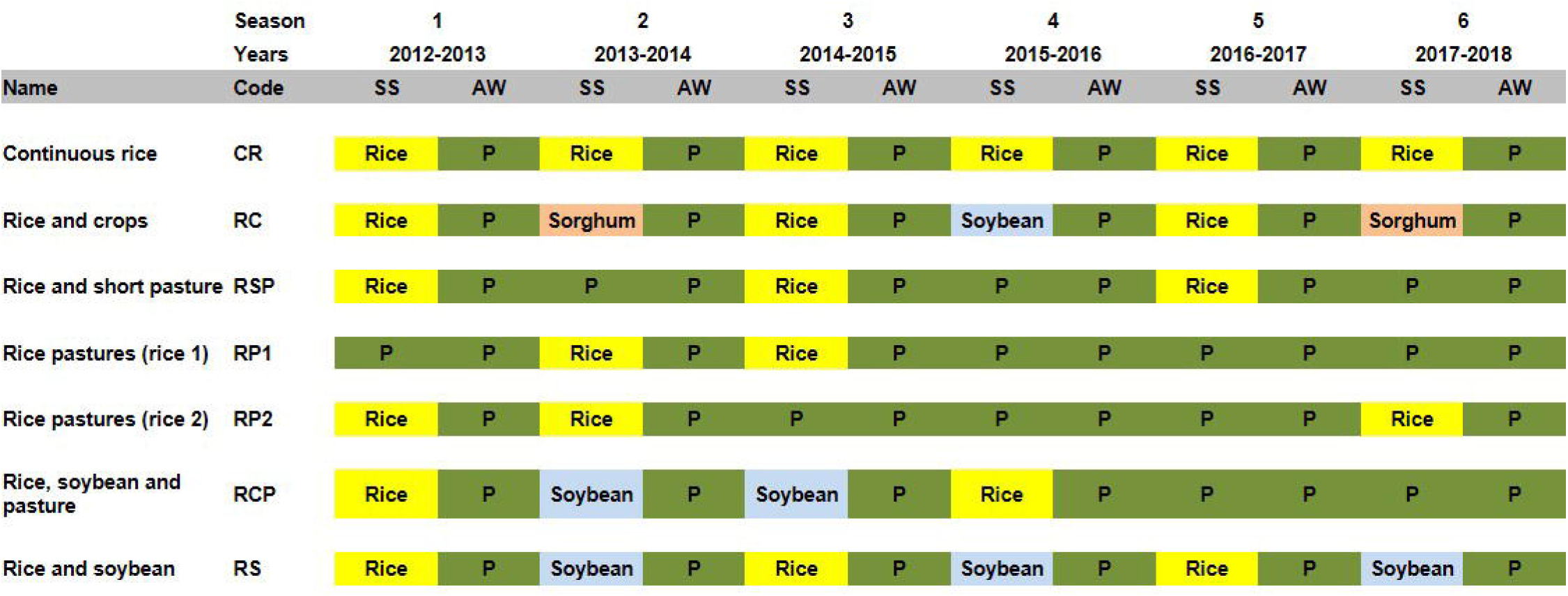
Scheme of the phases of the study site sampled at the Long-time experiment of Unidad Experimental de Paso de la Laguna, Treinta y Tres, Uruguay. Crop rotations, names and codes, and crop per season and year before soil sampling (SS= spring-summer, AW= autumn-winter, P= pasture).

The historical mean monthly temperature in the research area is 11.5±0.82 °C and 22.3±0.85 °C during winter and summer respectively; mean annual rainfall is 1360±315 mm; annual total potential evapotranspiration is 1138±177 mm. The mean annual precipitation (2012-2018) was 1290 mm, with a monthly mean of 107 mm for the period of the experiment, from September 2012 to September 2018.

Dominant soil type in the experiment was an Argialboll with a slope less than 0.5% (Durán et al., 2006). The soil (0-15 cm) was classified as clayey silt, containing 14.2 g kg^-1^ organic carbon, 0.14% total N, 28.8 mg exchangeable Mg kg^-1^, 659.6 mg available P kg^-1^, and 10.5 mg exchangeable K kg^-1^, and had a pH of 5.7 at the beginning of the experiment in September of 2012.

Management practices, fertilization, pesticide use, and irrigation were based on INIA Rice Program recommendations (Macedo et al., 2021).

### 2.2 Soil sampling

Soil samples were collected in September 2018 before rice seeding at the beginning of the second cycle of the experiment in seven different phases of the nine possible in the six rotations, including the two phases of the traditional rice and pasture system, before first and second rice (Fig. 1). In each 20 x 60 m plot, twenty 3-cm diameter soil cores were randomly taken from 0–10 cm depth of topsoil with a metal soil borer. The 20 soil cores were then mixed, homogenized, and separated into two samples. First, a 30-40-gram soil sample was air-dried under sterile conditions in Petri dishes, passed through a one mm metal sieve, transferred to 60 cm^3^ plastic jars and stored at –20 °C for DNA extraction. A second sample was air-dried and stored at 4 °C for soil chemical analysis.

### 2.3 Soil chemical analysis

The soil samples were analyzed for their chemical properties at the Laboratorio de Suelos, INIA La Estanzuela, Uruguay. Soil samples were air-dried at room temperature and passed through a one mm sieve to remove aboveground plant materials, roots, and stones. The pH was measured in triplicates in the supernatant of a 1:2.5 (w/v) soil:water solution ratio mixture after three minutes of shaking and 15 minutes of resting, using a pH meter according to Beretta et al. (2014). Available P content was measured by extraction and subsequent determination of the extracted P by absorbance with the Bray and Kurtz method (Bray and Kurtz, 1945). Soil organic carbon was measured by dry combustion of the sample and subsequent infrared detection of CO2 with the Wright and Bailey method (Wright and Bailey, 2001). Exchangeable macronutrients, such as magnesium and potassium, were measured by the ammonium acetate extraction method (1 M – pH 7,0), and reading the extract was completed by atomic emission (K) and atomic absorption (Mg) or by atomic emission with the ICP-OES team (Jackson, 1964).

### 2.4 DNA extraction, amplification, Illumina NovaSeq sequencing and data analysis

Genomic DNA was extracted from 1g of soil from 4 different samples from each sampled plot using the DNeasy PowerSoil Kit (Qiagen, Antwerp, Belgium) according with the manufacturer’s instructions. Concentration and quality of the DNA samples was measured by a NanoDrop ND-1000 spectrophotometer (Thermo Scientific, Wilmington, DE, USA). The four samples were combined and diluted to 20 ng/μl for final analysis. For bacteria, the V4–V5 region of the bacterial 16S rRNA gene was amplified by PCR using the primers set 515F (5’-GTGCCAGCMGCCGCGGTAA-3’) and 907R (5’-CCGTCAATTCCTTTGAGTTT-3’) and a high-fidelity polymerase. For fungal study, the V4 region of the 18S gene was amplified using the primers set 528F (5’-GCGGTAATTCCAGCTCCAA-3’) and 706R (5’-AATCCRAGAATTTCACCTCT-3’). This region of the 18S gene is more conserved than ITS or 28S ribosomal regions but can be used to estimate diversity at family level. The PCR conditions were 95°C for 5 min, followed by 25 cycles of denaturation at 95°C for 30 s, 50°C for 30 s, 72°C for 40 s, and a final extension period of 72°C for 7 min.

The PCR products were mixed with the same volume of 2× loading buffer and were subjected to 1.8% agarose gel electrophoresis for detection. Samples with a bright main band of approximately 450 bp were chosen and mixed in equal-density ratios. The obtained PCR products were purified using a GeneJET Gel Extraction Kit (Thermo Fisher Scientific, Waltham, MA, United States). Sequencing libraries were generated following manufacturer’s recommendations with NEB Next^®^ Ultra_™_ DNA Library Prep Kit for Illumina (NEB, USA). The constructed libraries were validated using an Agilent 2100 Bioanalyzer (Agilent Technologies, Palo Alto, CA, United States) and quantified with a Qubit 2.0 Fluorometer (Thermo Fisher). Finally, paired-end sequencing was conducted using an Illumina NovaSeq 6000 Sequencing System platform (Illumina, Inc., San Diego CA, United States) at CD Genomics, Co., Ltd. (NY, USA).

Paired-end reads obtained were merged using FLASH (Magoc and Salzberg, 2011). Quality filtering were then performed on the raw tags under specific filtering conditions of Trimmomatic (Bolger et al., 2014). Assembled sequences shorter than 300 bp in length were eliminated from the database. UCLUST in QIIME (Caporaso et al., 2010) was used to cluster the tags with 97% similarity and acquired the operational taxonomic units (OTUs). Singletons were removed from the datasets as recommended in previous studies (Lindahl et al. 2013). OTUs were then annotated using Silva taxonomic database.

### 2.5 Statistical analysis

Most statistical analyses were carried out by using the Vegan package (Oksanen et al., 2016) in R (version 4.0.1) (R Core Team, 2020) unless stated otherwise.

Differences between rotations were analyzed using a one-way analysis of variance (ANOVA), and an LSD test was performed for multiple comparisons using SAS version 9.4 (SAS Inc., Cary, NC, USA). Differences at *P* < 0.05 level were considered significant. The data reported for microbial diversity are mean values from the three replicates (plots) analyzed. Rarefaction curves and observed species were calculated using the Vegan package in R (Oksanen et al. 2019). The relative abundance-based coverage estimator (ACE), Chao1 lower-bound estimator of species richness (Chao1) richness index, and Shannon entropy (H’) and Simpson (S) diversity index were generated with estimateR function from the Vegan package in R (Oksanen et al. 2019). Alpha-diversity estimates for each diversity index between rice rotations were tested by one-way ANOVA and post-hoc Tukey’s HSD to identify significant differences among group means (P < 0.05). Non-metrical multidimensional scaling (NMDS) was run for comparison of beta-diversity measures on the Hellinger-transformed OTU tables, separately for bacterial and fungal communities. Ordination was run with parameters set on Bray-Curtis distance measure and two dimensions and total stress obtained in the Vegan package in R (Oksanen et al. 2019). Permutational multivariate analysis of variance (PERMANOVA, Anderson 2001) was used separately to determine the effect of the studied rotation systems compared with the traditional rice and pasture on the observed variance of the total bacterial and fungal communities. The bacterial and fungal communities were first tested for heteroscedasticity using betadisper function (*P* < 0.05) and permutational multivariate analysis of variance using adonis function in the Vegan package in R (Oksanen et al. 2019) on Bray-Curtis distance matrices obtained from OTU tables. Additionally, when a significant treatment effect was found a post hoc pairwise comparison was performed. Relative abundance of bacterial or fungal OTUs was analyzed for correlations to soil chemical properties using a canonical correspondence analysis (CCA; ter Braak, 1986) in R using Vegan and Spearman’s correlation coefficients. Analysis was also performed to assess possible relationships between relatively abundant bacterial and fungal phyla and soil chemical properties.

Bacterial and fungal taxa that characterize microbial communities in the different rice rotation soils studied were identified using linear discriminant analysis (LDA) effect size (LEfSe) (Segata et al., 2011). The classified taxa were discriminated accordingly in relation to having both high relative abundances within sampling soils and high frequencies across replicated plots within each rice rotation. LEfSe method identifies bacterial and fungal biomarkers via Kruskal-Wallis non-parametric test detecting significant taxa and performing an LDA effect size estimation. Biomarker taxa were identified with an α = 0.05 and an effect size threshold of LDA = 2. Biomarker taxa were then plotted across each rice rotation to help the visualization of the respective frequencies and relative abundances of occurrence (Segata et al., 2011). LEfSe analysis was performed through the Galaxy/Hutlab server (huttenhower.sph.harvard.edu/galaxy).

## 3. Results

### 3.1 Soil physico-chemical analysis

Among the chemical parameters analyzed, P and the relation C:N showed significant statistical differences between rice rotations (Table 1). Phosphorus concentration (*P* = 0.0162) was lower in continuous rice and the rotations with pastures in the system and higher in rotations of rice with crops (RC and RS). The relation C:N (*P* = 0.0415) was lower in the rotations with more time occupied with crops, rice, soybean, and/or sorghum in the sequence (Table 1). No statistical differences were found for K, Mg, organic C, N, organic matter, or pH among rotations after six years of the experiment (Table 1).

**Table 1.**
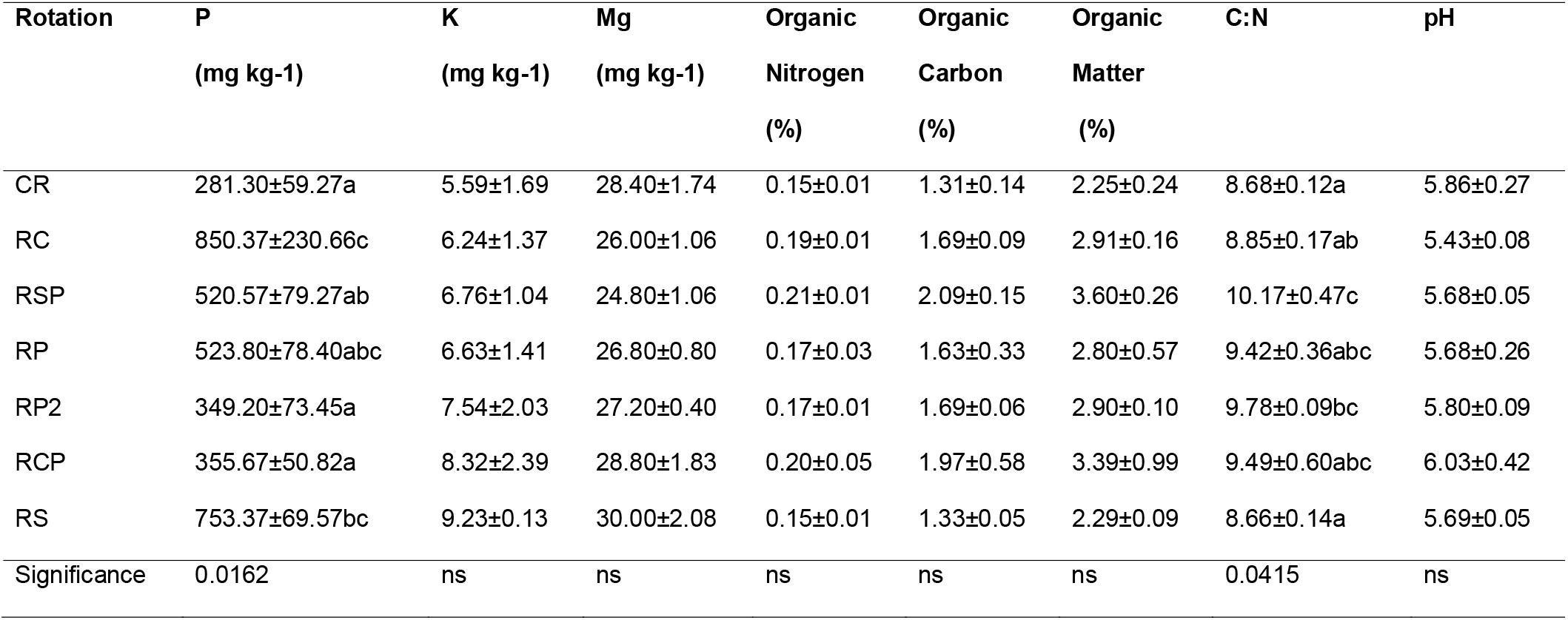
Soil chemical parameters evaluated for each of the rotations studied after six years at the Long-term experiment of Paso de la Laguna, Treinta y Tres, Uruguay. Rotations: CR= continuous rice, RC=rice and crops, RSP=rice and short pasture, RP= rice and pasture (after pasture), RP2= rice and pasture (after rice), RCP=rice, soybean and pasture, RS= rice and soybean. Values are means ± standard deviation of three replicates. Different letters indicate statistically significant differences between rotations (*P* < 0.05).

| Rotation | P<br>(mg kg<sup>-1</sup>) | K<br>(mg kg<sup>-1</sup>) | Mg<br>(mg kg<sup>-1</sup>) | Organic<br>Nitrogen<br>(%) | Organic<br>Carbon<br>(%) | Organic<br>Matter<br>(%) | C:N | pH |
| --- | --- | --- | --- | --- | --- | --- | --- | --- |
| CR | 281.30±59.27a | 5.59±1.69 | 28.40±1.74 | 0.15±0.01 | 1.31±0.14 | 2.25±0.24 | 8.68±0.12a | 5.86±0.27 |
| RC | 850.37±230.66c | 6.24±1.37 | 26.00±1.06 | 0.19±0.01 | 1.69±0.09 | 2.91±0.16 | 8.85±0.17ab | 5.43±0.08 |
| RSP | 520.57±79.27ab | 6.76±1.04 | 24.80±1.06 | 0.21±0.01 | 2.09±0.15 | 3.60±0.26 | 10.17±0.47c | 5.68±0.05 |
| RP | 523.80±78.40abc | 6.63±1.41 | 26.80±0.80 | 0.17±0.03 | 1.63±0.33 | 2.80±0.57 | 9.42±0.36abc | 5.68±0.26 |
| RP2 | 349.20±73.45a | 7.54±2.03 | 27.20±0.40 | 0.17±0.01 | 1.69±0.06 | 2.90±0.10 | 9.78±0.09bc | 5.80±0.09 |
| RCP | 355.67±50.82a | 8.32±2.39 | 28.80±1.83 | 0.20±0.05 | 1.97±0.58 | 3.39±0.99 | 9.49±0.60abc | 6.03±0.42 |
| RS | 753.37±69.57bc | 9.23±0.13 | 30.00±2.08 | 0.15±0.01 | 1.33±0.05 | 2.29±0.09 | 8.66±0.14a | 5.69±0.05 |
| Significance | 0.0162 | ns | ns | ns | ns | ns | 0.0415 | ns |

### 3.2 Sequencing information

A total of 1,557,096 paired-end sequences were obtained for 16S and 1,778,576 for 18S. Bacterial paired-end reads were overlapped to obtain high-quality tag sequences and after filtering 658,727 effective tag sequences were obtained with average lengths of 373 bp. After filtering, and discarding non-fungal tags, paired-end reads for 18S accounted for 647,806 effective tag sequences with an average length of 307 bp. OTU clusters were defined by a 97% identity threshold and 3662 and 807 OTUs were found for the bacterial and fungal communities, respectively.

### 3.3 Diversity and richness of the soil community

Obtained rarefaction curves showed that archaeal/bacterial communities were better represented than fungal communities (Fig. S2, Fig. S3). However, similar patterns were obtained for sequenced samples within both microbial groups from all soils suggesting that similar levels of diversity values were captured for each soil sampled (Fig. S2, Fig. S3).

Alpha diversity of the microbial communities in the soils studied was assessed by means of different richness and diversity indices. The observed number of bacterial OTUs was statistically different between rotations (*F* = 3.9577, *P* = 0.0159) with CR having the highest OTU number and the rotation RP2 and the agricultural rotations, RC and RS, having the lowest OTU numbers. Shannon index was statistically significant (*F* = 2.9044, *P* = 0.0470) with continuous rice different from the other rotations, except for RCP (Table 2). No significant differences were found in the richness indices, Chao1 and ACE, and Simpson diversity index among the different rotations studied (Table 2).

**Table 2.** Diversity indices at the OTU level of prokaryotic communities (archaea/bacteria) for each rotation studied. Rotations: CR = continuous rice, RC = rice and crops, RSP = rice and short pasture, RP = rice and pasture (after pasture), RP2 = rice and pasture (after rice), RCP = rice, soybean and pasture, RS = rice and soybean. Values are means ± standard deviation. Different letters indicate statistically significant differences between rotations (*P* < 0.05).

|  | **CR** | **RC** | **RSP** | **RP** | **RP2** | **RCP** | **RS** |
| --- | --- | --- | --- | --- | --- | --- | --- |
| Observed species | 1814±31a | 1658±28b | 1780±25ab | 1683±65ab | 1726±21ab | 1753±15ab | 1704±34ab |
| Shannon | 5.839±0.05 | 5.634±0.04 | 5.699±0.014 | 5.702±0.03 | 5.683±0.06 | 5.786±0.04 | 5.690±0.04 |
| Simpson | 0.991±0.0001 | 0.989±0.0001 | 0.989±0.001 | 0.990±0.001 | 0.990±0.001 | 0.991±0.001 | 0.989±0.001 |
| Chao1 | 2189±18 | 2073±9 | 2188±7 | 2115±66 | 2115±9 | 2118±34 | 2153±44 |
| ACE | 2218±37 | 2115±20 | 2246±13 | 2142±64 | 2168±23 | 2162±29 | 2147±38 |

Fungal OTU numbers were higher in rotation RP2 and at lower OTU number was found in RCP. Shannon diversity index varied between rotations (*F* = 3.0342, *P* = 0.0408), with the highest value in rotations RS and RP2 and lowest value in CR and RP (Table 3). No significant differences for ACE and Chao1 richness indices and Simpson diversity index (*P* = 0.07) for fungal communities were observed among rice rotation soils (Table 3).

**Table 3.** Diversity indices at the OTU level of fungal communities for each rotation studied. Rotations: CR = continuous rice, RC = rice and crops, RSP = rice and short pasture, RP = rice and pasture (after pasture), RP2 = rice and pasture (after rice), RCP = rice, soybean and pasture, RS = rice and soybean. Values are means ± standard deviation. Different letters indicate statistically significant differences between rotations (*P* < 0.05).

|  | CR | RC | RSP | RP | RP2 | RCP | RS |
| --- | --- | --- | --- | --- | --- | --- | --- |
| Observed_species | 292±12 | 322±30 | 329±9 | 321±15 | 343±13 | 298±36 | 299±21 |
| Shannon | 2.69±0.17b | 2.99±0.08a | 2.98±0.05a | 2.63±0.10b | 3.03±0.10a | 2.83±0.08ab | 3.06±0.04a |
| Simpson | 0.826±0.049 | 0.891±0.012 | 0.896±0.002 | 0.836±0.010 | 0.900±0.013 | 0.875±0.013 | 0.893±0.007 |
| Chao1 | 401±22 | 441±26 | 427±8 | 423±25 | 426±9 | 399±38 | 399±8 |
| ACE | 407±31 | 417±45 | 432±11 | 426±16 | 426±12 | 419±24 | 378±18 |

### 3.4 Archaeal, bacterial and fungal community composition

The prokaryote community was represented at most relative abundance by domain Bacteria (98.9%) not Archaea (1.02%). Whole bacterial community composition within the different soils studied showed *Proteobacteria*, *Firmicutes*, *Chloroflexi*, *Actinobacteria*, *Planctomycetes* and *Acidobacteria* as the most OTU rich at the phylum level (Fig. S4). The bacterial phyla with the highest relative abundance in the whole study were *Actinobacteria* (35.8%), *Firmicutes* (19.9%) and *Proteobacteria* (17.4%). The remaining important phyla (*Acidobacteria*, *Chloroflexi*, *Verrucomicrobia*, *Planctomycetes* and others) accounted for less than 10% but more than 1% of the abundance each. Sequences obtained for each rotation were associated with nine bacterial phyla with >1% of relative abundance in at least one rotation or 18 phyla with >0.1% of relative abundance (Fig. 2). *Actinobacteria*, *Firmicutes*, and *Proteobacteria* ranged from 32.5–-39.6%, 18.3–-22.4%, and 16–-19.1% relative abundance, respectively, according to the rotation (Fig. 2). *Chloroflexi*, *Acidobacteria*, *Verrucomicrobia* and *Planctomycetes* were found at relative abundances <10% but >1%. Relative abundances >1% were found for *Euryarchaeota* (Archaea) and *Bacterioidetes* only in CR and in CR and RSP, respectively (Fig. 2). Relative abundance of *Actinobacteria* did not differ between the seven rotations studied (*F* = 1.0003, *P* = 0.4626), but the lowest relative abundances were found in the rotations with 100% of agriculture of rice with other crops (RC and RS). Continuous rice incremented the abundance of *Proteobacteria* (*F* = 3.6504, *P* = 0.0215), and the relative abundance was significantly higher than in the rotations including other crops like soybean and sorghum (rotations RC, RCP and RS), but not different from rotations with more pastures in the cycle (Fig. 3). *Chloroflexi* was incremented in treatments with more crops in the cycle, but differences were not statistically significant (*F* = 2.1706, *P* = 0.1089). Comparing the rotations by pairs showed CR differed statistically from the less intensive rotations of rice with longer pastures cycles (rotations RP, RP2 and RCP). *Verrucomicrobia* differed statistically between rotations (*F* = 3.7177, *P* = 0.0201) with higher relative abundances in the rotations of RS and second rice with long pastures and RC (Fig. 3). Other less represented phyla of bacteria or archaea were statistically different within treatments. *Crenarchaeota* (Archaea, *F* = 15.1422, *P* = 0.0001) and *Nitrospirae* (*F* = 3.5084, *P* = 0.0248) were more abundant in continuous rice than other rotations, while *Cyanobacteria* (*F* = 4.6344, *P* = 0.0085) was more abundant in continuous rice than other rotations, except for RSP, and the rotations with other crops had the lowest relative values (Fig. S5).

**Fig.2.**
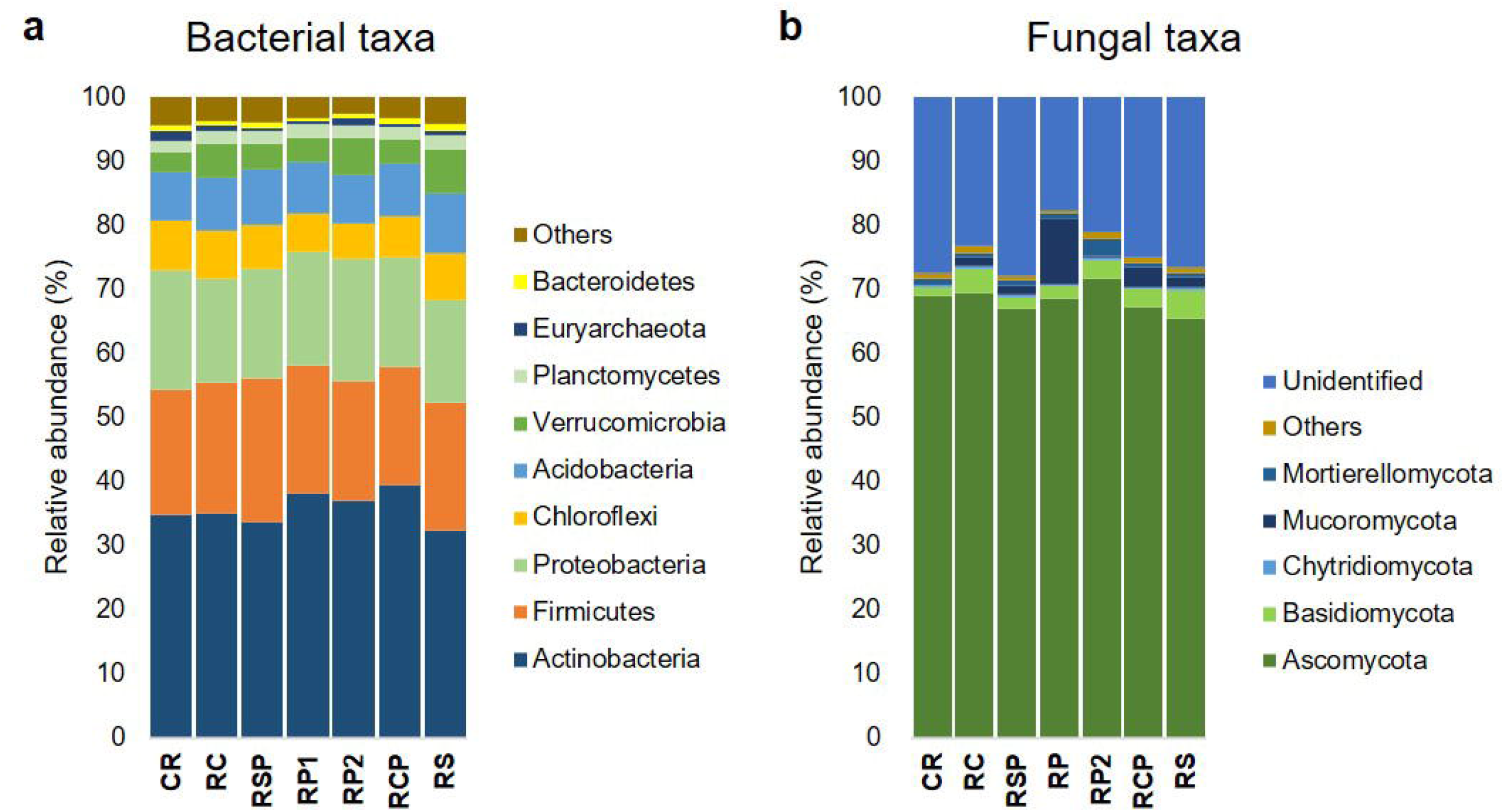
Relative abundance found of a) bacterial phyla, and b) fungal classes according with rotation studied. Rotations: CR=continuous rice, RC=rice and crops, RSP=rice and short pasture, RP=rice and pasture (after pasture), RP2=rice and pasture (after rice), RCP=rice, crops and pasture, and RS=rice and soybean (see Fig. 1).

**Fig. 3.**
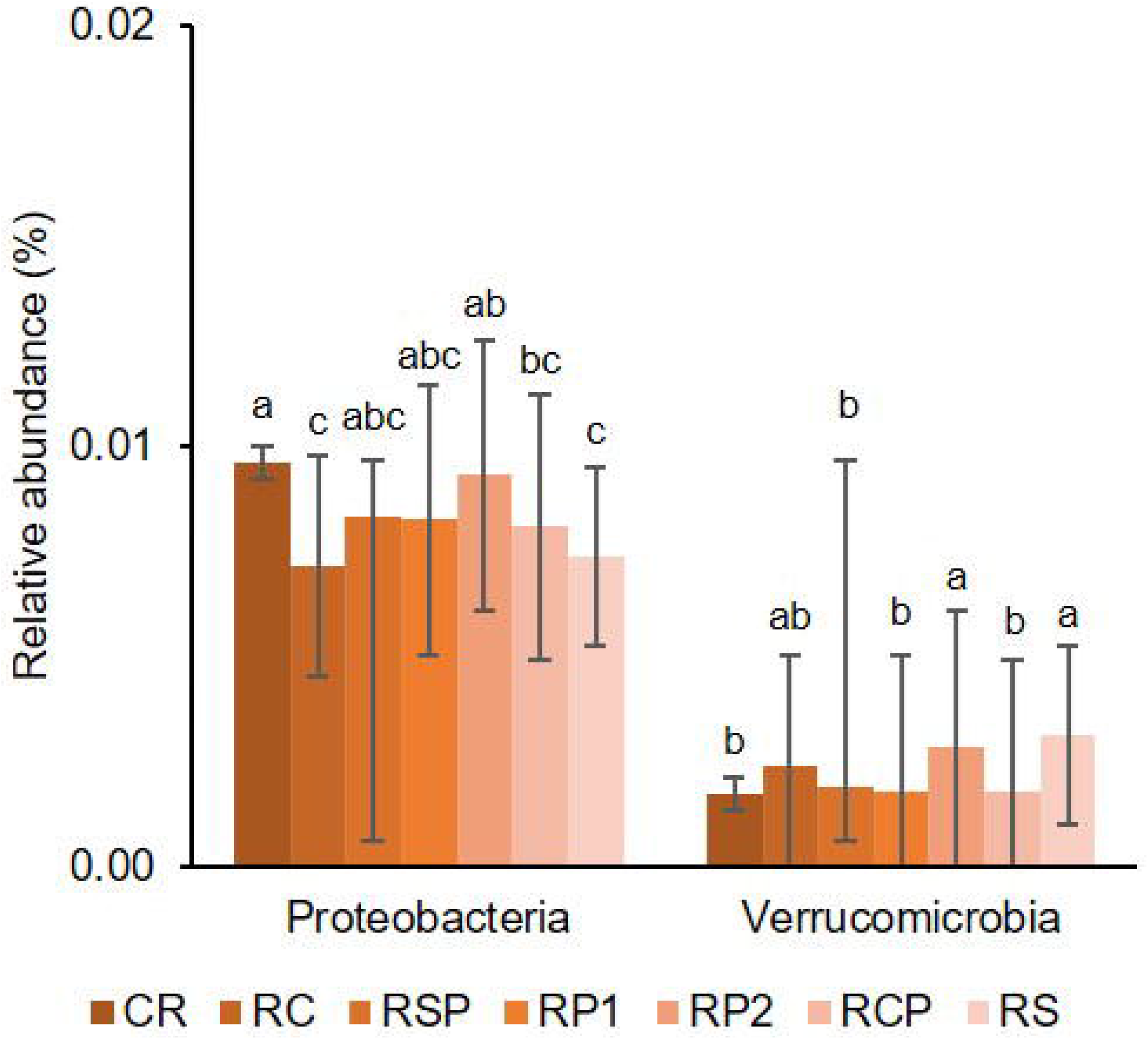
Relative abundance found of bacterial phyla *Proteobacteria* and *Verucomicrobia* according with rotation studied. Rotations: CR=continuous rice, RC=rice and crops, RSP=rice and short pasture, RP=rice and pasture (after pasture), RP2=rice and pasture (after rice), RCP=rice, crops and pasture, and RS=rice and soybean (see Fig. 1). Bars represents standard errors based in three replications and different letters above each bacterial phylum indicate significant differences (*P*<0.05) according with Fisher’s LSD.

Fungal community composition in the soils from different rotations showed that most OTUs (388) remain unidentified (Fig. S4). *Chytridiomycota* (93), *Ascomycota* (84), *Cryptomycota* (58), *Basidiomycota*, (40) and *Aphelidea* (39) were the most OTU rich phyla in the kingdom Fungi (Fig. S4). The highest relative abundance was found for *Ascomycota* (68.4%), ranging from 65.5% in RS rotation to 71.8% in RP2 (Fig. 2). *Basidiomycota* (2.71%), *Mucoromycota* (2.57%), and *Mortierellomycota* (0.987%), were fungal taxa accounting for relative abundances around 1% or more. The phyla, *Aphelidea*, *Cryptomycota*, *Mucoromycota*, *Neocallimastigomycota* and *Incertae sedis* (Fungi) were represented by relative abundances of <1% but over 0.1%. Other taxa were represented but with very low relative abundances (<0.1%, including *Blastocladiomycota*, *Glomeromycota*, *Nuclearida*, *Olpidiomycota*, and *Zoophagomycota*). Some of these phyla varied significatively among rotations, including *Cryptomycota* (*F* = 3.1742, *P* = 0.0351), *Glomeromycota* (*F* = 2.9461, *P* = 0.0449), and *Incertae sedis* (*F* = 3.8275, *P* = 0.0181) (Fig. 4). Within the *Ascomycota*, the most abundant fungal phyla, *Sordariomycetes*, *Eurotiomycetes* and *Dothideomycetes*, accounted for >95% of the relative abundance. Orders with the highest relative abundances were *Hypocreales* (30.8%), *Eurotiales* (25.5%), *Pleosporales* (19%), *Sordariales* (16.3%), *Capnodiales* (2.4%) and *Boliniales* (1.3%). Others, unidentified (2.3%), and *Incertae sedis* (2.4%) completed the list of fungal taxa found. *Basidiomycota, Tremellales* (57.8%), *Atheliales* (20.7%), *Ceratobasidiaceae* (5.9%), *Cystofilobasidiaceae* (3.9%) and *Sporidiobolaceae* (2.9%) were the most abundant taxa within the phylum. *Tremellales* was represented exclusively by *Saitozyma* spp. and *Atheliales* exclusively by *Athelia rolfsii* (Table S2). Four fungal taxa (*Fusarium oxysporum*, unidentified *Aspergillaceae*, *Chaetomium globosum* and unidentified Fungi) accounted for >50% of the relative abundance observed (Table S2).

**Fig. 4.**
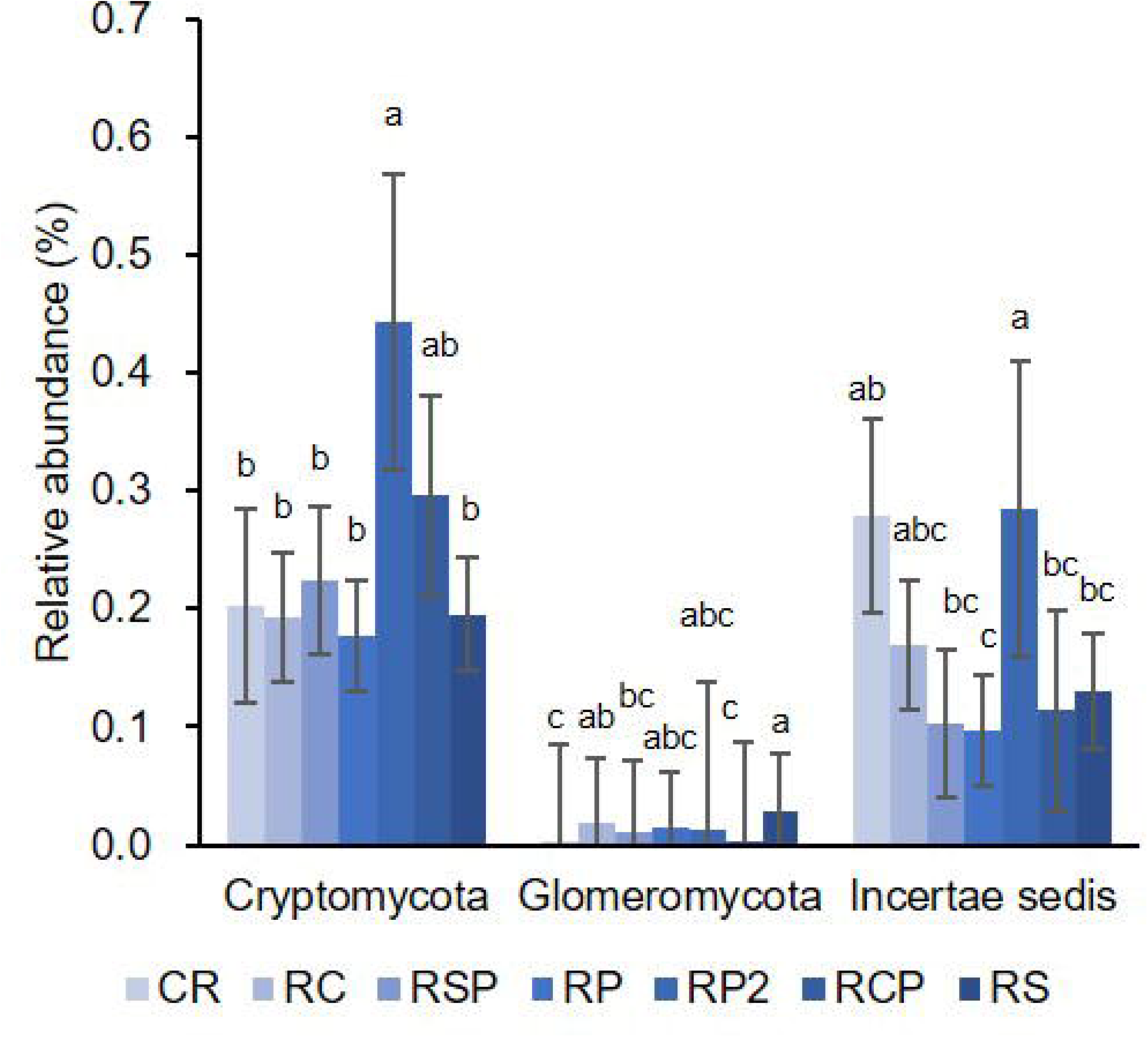
Relative abundance found of fungal classes *Cryptomycota*, *Glomeromycota* and *Incertae sedis* according with rotation studied. Rotations: CR=continuous rice, RC=rice and crops, RSP=rice and short pasture, RP=rice and pasture (after pasture), RP2=rice and pasture (after rice), RCP=rice, crops and pasture, and RS=rice and soybean (see Fig. 1). Bars represents standard errors based in three replications and different letters above each fungal class indicate significant differences (*P*<0.05) according with Fisher’s LSD.

### 3.5 Microbial community composition

Non-metric multidimensional scaling (NMDS) ordination constructed on total bacterial OTU composition showed that samples clustered in three groups consistent with the percentage of years occupied with rice in the rotation cycle. One group formed alone with rotation CR (100% rice); a second group formed with the cluster of rotations with other crops or short pasture (RC, RS, and RSP with 50% of time occupied with rice); and a third group formed with more lax rice rotations, including traditional rice and pastures (RP, RP2, and RCP 33–-40% of time occupied with rice, Fig. 5). NMDS ordination for archaeal/bacterial communities resulted in a two-dimensional final resolution with a stress value of 0.196 (Fig. 5). PERMANOVA indicated a significant effect of the rotation system upon the bacterial OTU assemblages (*F* = 1.548, *P* = 0.0012). Permutation test for homogeneity of multivariate dispersions showed a nonsignificant dispersion (*F* = 0.223, *P* = 0.963). However, post hoc analysis failed to discriminate differences between pairs of soils (*P* < 0.05).

**Fig. 5.**
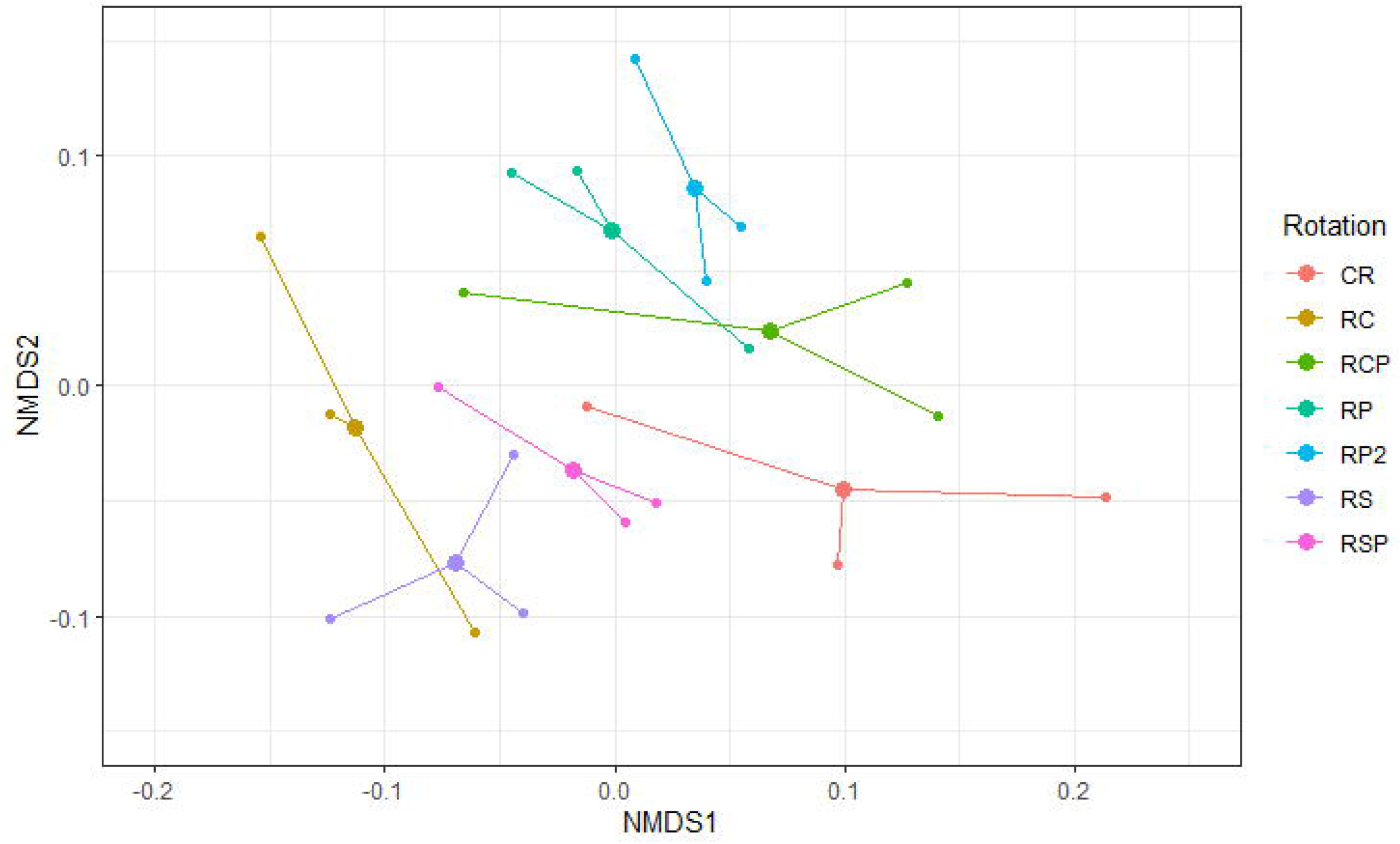
Non-metric multidimensional scaling (NMDS) ordination based on Bray-Curtis distance for differences in composition of prokaryotic communities in the different rotations studied (see Fig. 1). Rotations: CR=continuous rice, RC=rice and crops, RSP=rice and short pasture, RP=rice and pasture (after pasture), RP2=rice and pasture (after rice), RCP=rice, crops and pasture, and RS=rice and soybean (Stress: 0.196).

The NMDS analysis of fungal communities showed a separation of communities according to rotation, but without grouping within different soils. The analysis showed a separation into three groups, mostly according to the previous crop/pasture as a rice antecessor. These groups consisted of two rotations with rice as antecessor (CR and RP2), two rotations with soybean/sorghum as antecessors (RC and RS) and a third group with one to three years of pastures as antecessors (RP, RSP and RCP, Fig. 6). The NMDS ordination of fungal communities also resulted in a two-dimensional final resolution with a stress value of 0.16 (Fig. 6). PERMANOVA indicated a significant effect of the rotation system upon the fungal OTU assemblages (*F* = 1.635, *P* =0.0051). Permutation test for homogeneity of multivariate dispersions showed a nonsignificant dispersion (*F* = 0.587, *P* = 0.736). However, post hoc analysis failed to discriminate differences between pairs of soils (*P* < 0.05).

**Fig. 6.**
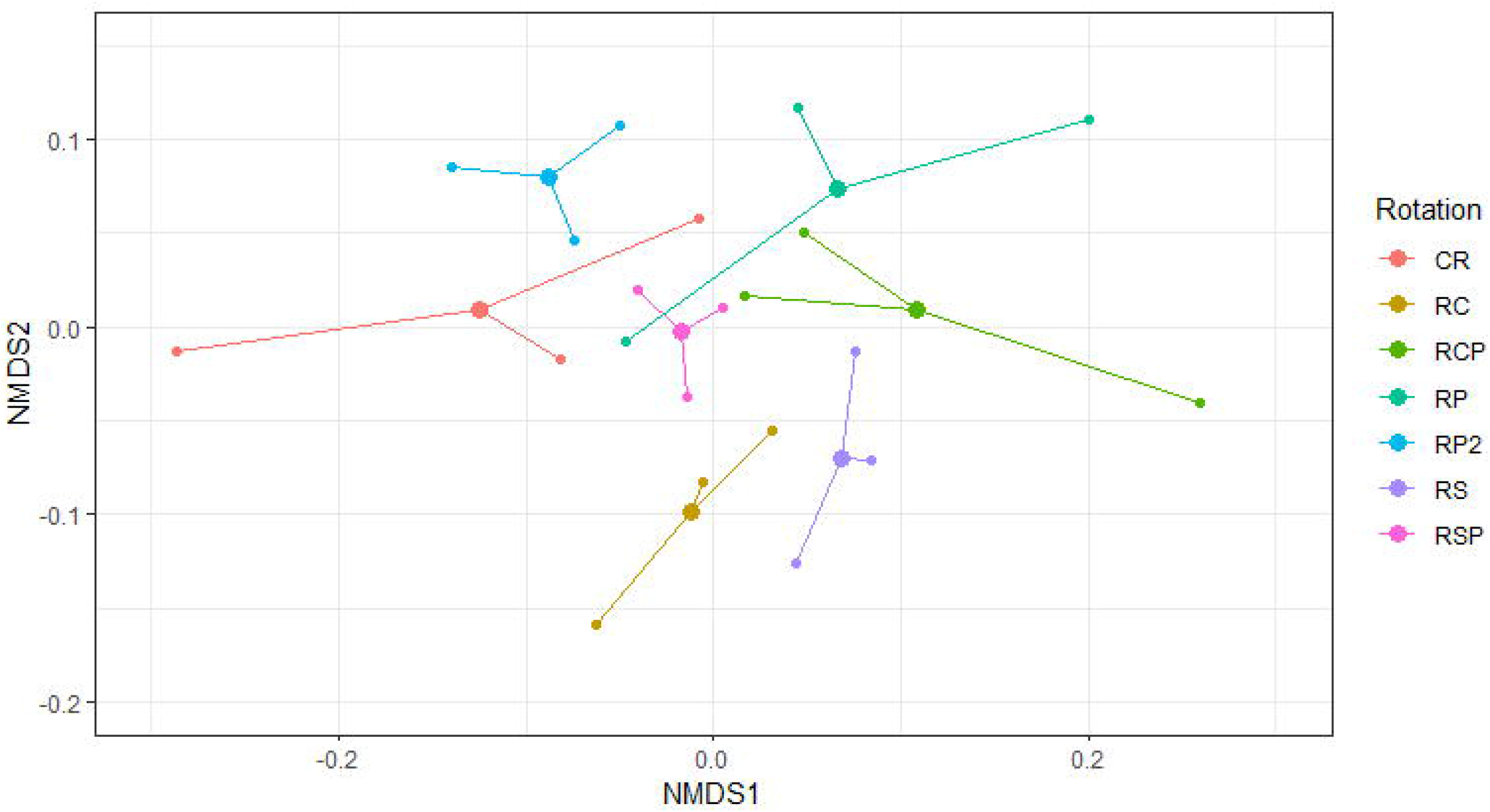
Non-metric multidimensional scaling (NMDS) ordination based on Bray-Curtis distance for differences in composition of fungal communities in the different rotations studied (see Fig. 1). Rotations: CR=continuous rice, RC=rice and crops, RSP=rice and short pasture, RP=rice and pasture (after pasture), RP2=rice and pasture (after rice), RCP=rice, crops and pasture, and RS=rice and soybean (Stress: 0.16).

Canonical correspondence analysis (CCA) was used to determine the correlation of soil chemical results on individual archaeal/bacterial and fungal OTUs (Fig. 7, Fig. 8). Individual bacterial OTUs appear to be weakly influenced by P, pH, and Mg and fungal OTUs appear to be weakly influenced by P, as shown by the OTUs dispersion around the vectors representing chemical values on the CCA (Fig. 7, Fig. 8). Additionally, rho values of Spearman correlations of the most abundant (> 0.01%) bacterial phyla with soil chemical parameters were analyzed. Seven of the bacterial phyla (*Actinobacteria* (K), *Proteobacteria* (P), *Verrucomicrobia* (P), *Synergistetes* (C and N), *Thaumarchaeota* (pH), *Cyanobacteria* (K) and *Nitrospirae* (C/N)) significantly (*P* < 0.05) correlated with at least one of these chemical parameters. Also, nine phyla (*Actinobacteria* (P), *Proteobacteria* (K), *Verrucomicrobia* (K), *Synergistetes* (K), *Thaumarchaeota* (P), *Planctomycetes* (K), *Chloroflexi* (C:N), *Spirochaetes* (C and N) and *Gemmatimonadetes* (P)) were weakly (*P* < 0.10) correlated with at least one chemical parameter (Table S3). Within fungal phyla, rho values of Spearman correlations of fungal classes showed that five (*Ascomycota* (K and Mg), *Chytridiomycota* (P), *Glomeromycota* (P), *Nuclearida* (P) and *Zoophagomycota* (P)) were correlated (*P* < 0.05) with one or two chemical parameters evaluated. Also, five fungal phyla (*Aphelidea* (N, pH), *Chytridiomycota* (Mg), *Cryptomycota* (C/N), *Glomeromycota* (pH), *Neocallimastigomycota* (pH)) were weakly (*P* < 0.10) correlated with at least one chemical parameter (Table S3).

**Fig. 7.**
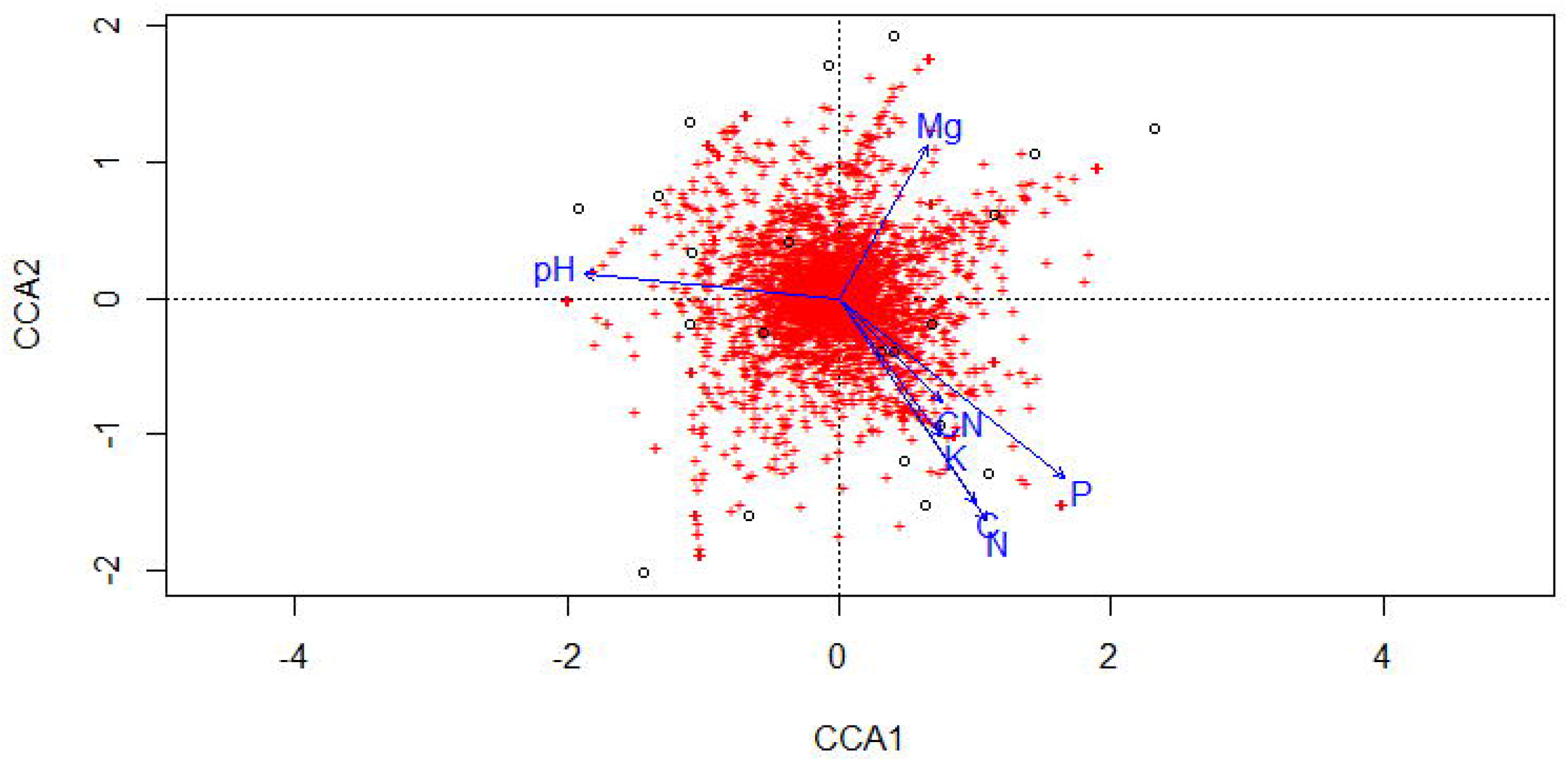
Canonical correspondence analysis (CCA) of bacterial OTUs in relation to measured soil physico-chemical parameters.

**Fig. 8.**
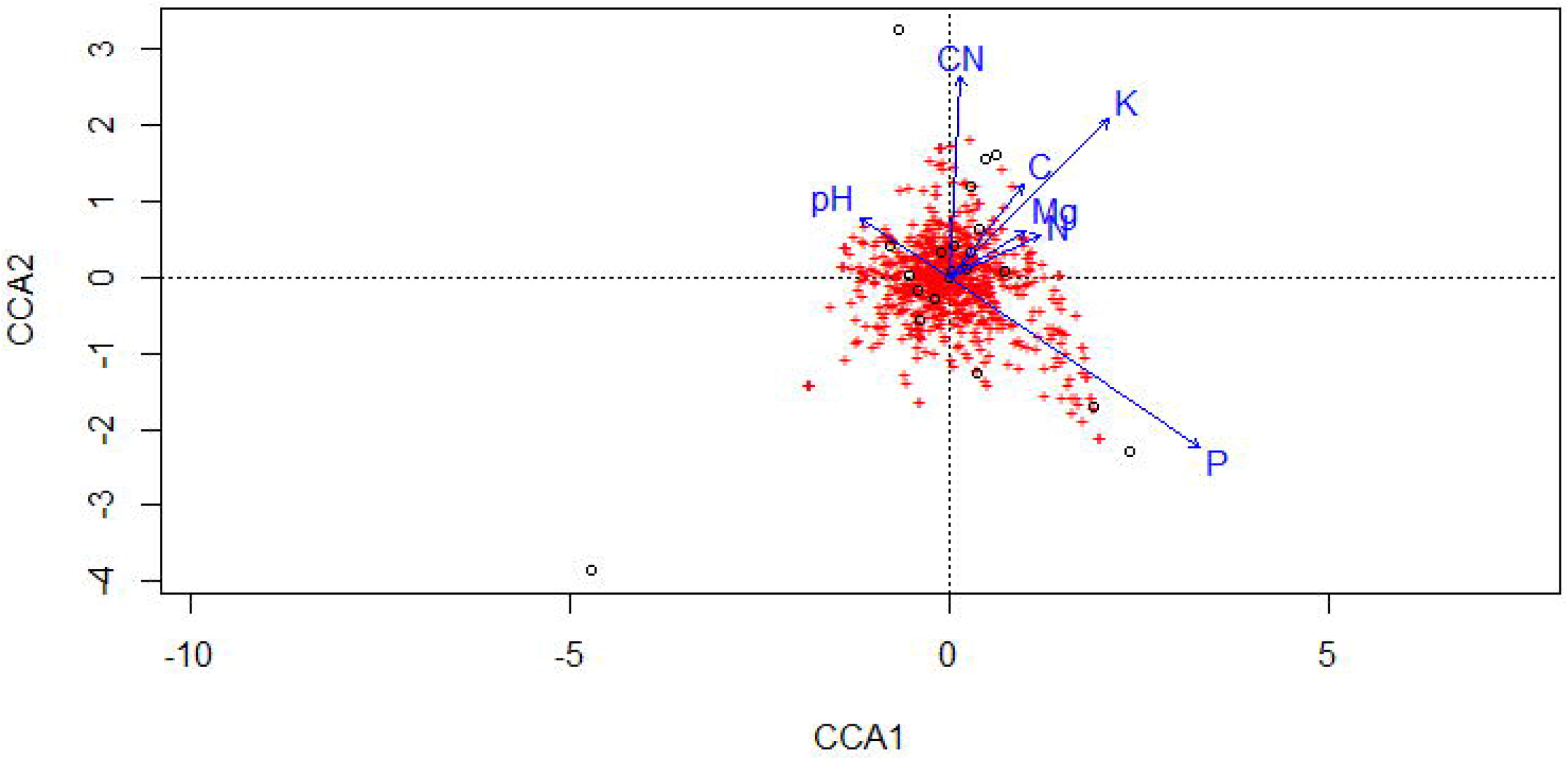
Canonical correspondence analysis (CCA) of fungal OTUs in relation to measured soil physico-chemical parameters.

Taxonomic biomarkers for archaea/bacteria and fungi among rice rotations were detected via Linear discriminant analysis Effect Size (LEfSe). The LEfSe analysis for bacterial communities revealed that there were more biomarkers with larger effect sizes (LDA score > 2.5) associated with the CR rotation, while agricultural rotations showed the lowest number of bacterial biomarkers (Fig. 9, Fig. S6). Rotations including pastures showed intermediate numbers of biomarker taxa (LDA score > 2.5). Considering the traditional rice and pasture rotation, prokaryotic indicator taxa that contributed the most were *Acetobacteraceae* and *Promicromonosporaceae* after pasture (RP), while *Betaproteobacteriales* and *Pseudonocardia* characterized the rotation after rice (RP2) (Fig. 9, Fig. S6). Other rotations after pasture that showed intermediate numbers of biomarkers (LDA score > 2.5) include *Pseudonocardiales* and *Actinomycetospora* as the most characteristic for RCP and *Ruminococcaceae, Paenibacillaceae* and *Ammoniphilus* as the most characteristic for RSP (Fig. 9, Fig. S6). Biomarker taxa for CR included Bacteria *Anaerolineae, Methylocystis, Nakamurella, Mycobacterium*, uncultured *Chloroflexi*, and *Phenylobacterium* and Archaea *Euryarchaeota*, *Bathyarchaeia, Chrenarchaeota* and *Methanosarcina* (Fig. 9, Fig. S6). More agricultural rotations were characterized by *Ktedonobacterales* for RC and *Rokubacteriales* for RS (Fig. 9, Fig. S6). Fungi contributed less biomarkers for rotation characterization and only eight taxa were found with significative values (LDA score > 2.5) for four rotations, CR, RP, RP2, and RS. Interestingly, rice and pasture, after pasture or rice (RP and RP2), were characterized by five biomarker taxa, but these were unidentifiable at the taxonomic level, except for an *Acremonium* sp. (Fig. S7).

**Fig. 9.**
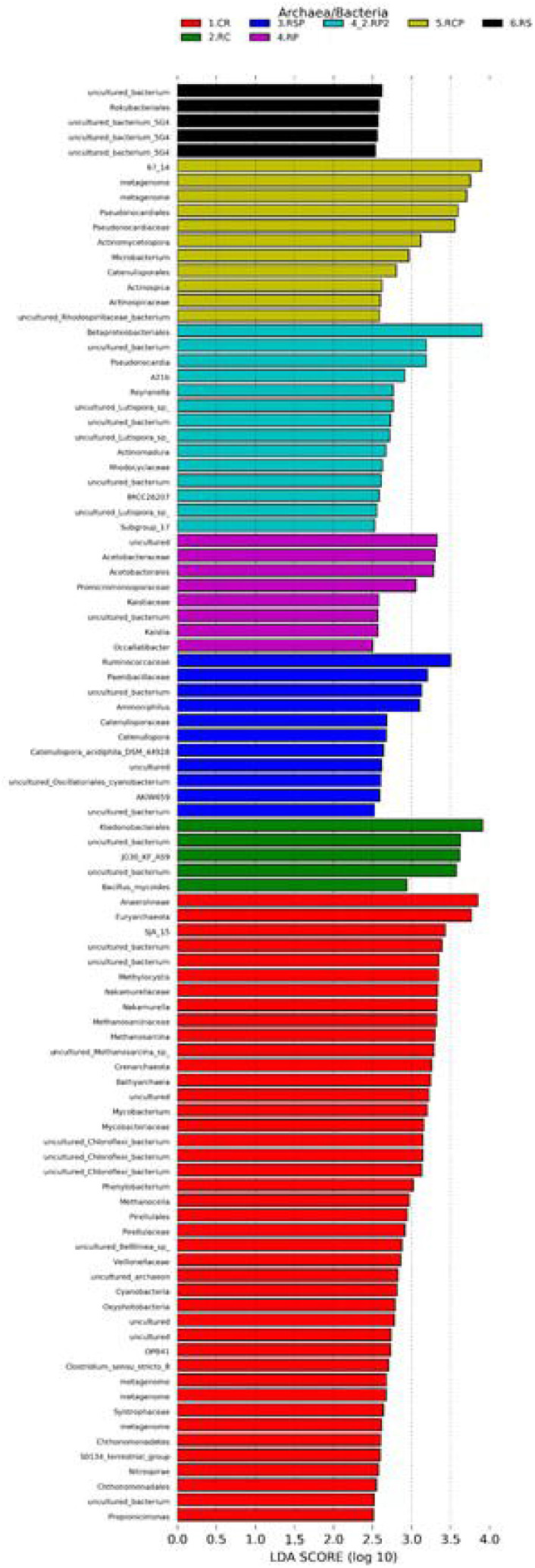
Linear discriminant analysis Effect Size (LEfSe) (log10 LDA score) of prokaryotic (Archaea/Bacteria) OTUs, which most likely explain differences between rice rotations after a complete cycle of six years. Archaea/Bacteria OTUs are classified at the highest resolvable taxonomic level. Rotations: CR= continuous rice, RC=rice and crops, RSP=rice and short pasture, RP= rice and pasture (after pasture), RP2= rice and pasture (after rice), RCP=rice, soybean and pasture, RS= rice and soybean.

## 4. Discussion

Metagenome study results based on NovaSeq sequencing from the first rotational cycle of the LTE demonstrated that differences in crops and management practices shaped soil microbial communities after six years, although at different degrees depending on the group analyzed.

Physicochemical analysis show that only P and C:N varied among the rotations studied (Table 1). Phosphorous concentration was higher in rotations of rice associated with other crops, like soybean and sorghum, due to the residual P from the previous fertilization. The C:N was higher in rotations associated with pastures previous to rice being one to three years long in the rotation due to the incorporation of carbon by the residues from the pastures. No clear correlations were obtained between different groups of prokaryotes and fungi and the soil parameters, except for some such as P and C:N (Table S3), and this could be due to the few changes occurring in the soils after six years of the experiment.

Differences among bacterial and fungal richness were found between rotations studied; the highest bacterial diversity and lowest fungal diversity was found in CR (Table 2 and 3). The high bacterial diversity in CR is contrary to reports indicating a decrease in diversity of microorganisms in simplified rotations of agricultural systems (Xuan et al. 2012; Venter et al. 2016). However, except for the OTU number no statistical differences in abundance and diversity index were found between rotations after six years (Table 1). The most abundant bacterial phyla were *Actinobacteria, Firmicutes*, *Proteobacteria*, and to a lesser extent (< 10% relative abundance) *Acidobacteria*, *Chloroflexi*, *Verrucomicrobia* and *Planctomycetes*. These dominant bacterial phyla were previously reported as ubiquitous in worldwide soils (Delgado-Baquerizo et al. 2018). Although *Actinobacteria* and *Firmicutes* are not commonly reported as the most abundant in rice soils, these were the most abundant phyla found, respectively, in experiments of preincubated and stored drained and flooded paddy soils (Wang et al. 2015). *Actinobacteria* sequences ranged from 32.5% (RS) to 39.6% (RCP) of relative abundance, quite higher than previous reports for this group in paddy soils, which ranged from 10 to 16% (Fernández et al. 2013; Jiang et al. 2016; Chen et al. 2016; Ji et al. 2020) or as low as <3% (Xuan et al. 2012) but slightly above the world relative abundance reported previously for this group (>30%, Delgado-Baquerizo et al. 2018). *Actinobacteria* abundances were higher in rotations with pasture and lower with crops, but without statistical differences, perhaps because only one cycle was completed, and more years are needed to differentiate the communities. Results are consistent with Lauber et al. (2008) who found more abundant *Actinobacteria*, also not significantly, in pasture soils, and Araujo et al. (2020) who reported reduced actinobacterial diversity with agriculture-based disturbance. *Actinobacteria* are Gram positive, aerobic, saprophytic bacteria reported across a diverse range of ecologies and environments, with higher abundances in warm dry soils due to its ability to tolerate drought (Delgado-Baquerizo et al. 2018; Ren at al. 2018; Araujo et al. 2020). Some members are known for their ability to secrete diverse types of metabolites such as antibiotics and extracellular enzymes (Selvakumar et al. 2014). The relatively high abundance of *Actinobacteria* found could be influenced by the timing of the soil sampling before rice sowing. The stress imposed during the fallow time after the use of herbicides in the previous pasture or cover crop established conditions of dry soil with a high contribution of organic residues that favored this group (Selvakumar et al. 2014; Araujo et al. 2020). *Firmicutes* was the bacterial phylum with the second highest relative abundance (19.9%) but without differences between rotations. The abundance is not surprising since it was the most abundant phylum found in a previous study from rice and pastures soil (28-40%, Fernández et al. 2013) and grasslands (17%, Garaycochea et al. 2020) in Uruguay. Data from paddy soils indicate that *Firmicutes* was represented by low percentages (<10%) in most situations (Xuan et al. 2012; Jiang et al. 2016; Ji et al. 2020). Also, the two most common, and four of the twenty most common bacterial OTU were *Firmicutes* members, mainly *Bacillales*, accounting for >10% of the total abundance (Table S1). *Firmicutes* is the most important group fermenting organic matter in anoxic conditions and has the ability to form endospores adapting to the stress of the dry conditions imposed by the fallow in the transitions of the pasture phase of the rotation (Fernández et al. 2013). *Proteobacteria* accounted for 17.4% of the sequences, ranging from 16% in rotations with crops (RC and RS) to 19% (RP2). These relative abundances are quite lower than world reported mean (>35%, Delgado-Baquerizo et al. 2018), and slightly lower than values reported in rice fields in Uruguay (20-28%, Fernández et al. 2013) and worldwide (26-38%, Jiang et al. 2016; Chen et al. 2016; Zhao et al. 2017). The abundance of *Proteobacteria* decreased according to the type of crop in the rotation and not according to the level of intensification. Higher values were found in rice with pastures (RP, RP2 and RSP) and CR and lower in rotations with other crops such as soybean and sorghum (RC, RCP, RS). Furthermore, *Proteobacteria* was the main or second most important phyla found in prokaryotic communities of Uruguayan soils in grassland and rice-pasture rotation (Fernández et al. 2013; Garaycochea et al. 2020). Members of *Proteobacteria* play important roles such as P solubilization, N fixation, plant growth promotion, and indication of copiotrophic soils, suggesting a better nutrient status and associated with higher rice yields (Huang et al. 2020).

Fungal communities were represented mainly by unidentified taxa (388 OTUs) that show the complexity of this group of organisms and denote the limited information available on the 18S region. More studies are needed that allow a more efficient identification at a lower taxa level, for example, by sequencing other regions such as ITS. Within the identified groups, *Ascomycota* dominated in all the soils in accordance with the fact that this is the most abundant group of fungi on a world scale with an average >68% relative abundance (Egidi et al. 2019). *Sordariomycetes* was the most abundant class represented, which includes *Pyricularia oryzae* (causal agent of rice blast) and *Nakataea oryzae* (causal agent of stem rot of rice), the two most devastating pathogens of rice in Uruguay (Martínez 2016; Martínez et al. 2018). *Hypocreales* was the most represented order of fungi and this was mostly represented by *Bionectriaceae* and *Nectriaceae* species, such as *Fusarium oxysporum* (Table S2). *Basidiomycota* and *Mucoromycota* were the second most abundant phyla with similar abundances (2.71% and 2.57% respectively). *Basidiomycota* was most represented in agricultural rotations (RC and RS, Fig. 2) with most sequences assigned to *Athelia rolfsii* (Table S2) a common pathogen of soybean in Uruguay (Stewart and Rodríguez, 2016). *Mucoromycotina* was highly represented only in one of the samples in the RP rotation associated with a high sequence count of the saprophytic *Rhizopus oryzae* (Table S2). Only *Cryptomycota*, *Glomeromycota* and *Incertae sedis* had differences in abundance between rice rotations (Fig. 4). Although *Glomeromycota* (also known as arbuscular mycorrhizal fungi or AMF) abundances were very low, higher values were found in rotations with soybean (RS and RC, Fig. 4), as reported in the same experiment by Rodríguez-Blanco and Giménez (2019) who found that roots of soybean plants were heavily colonized by *Glomeromycota* communities. However, more studies are needed on the relationship of *Glomeromycota* with the rice and pastures rotations (RP and RP2) since no differences in abundances were found with soybean rotations (Fig. 4).

These results demonstrated the responses of the bacterial/archaea and fungal communities to crop rotation and intensification despite the similarity between some of the results and the fact that only one cycle elapsed. Comparing the total OTUs of the bacterial communities showed that the rotations were clustered into three groups. The lax rotations with 33–40% of time with rice (with two to three years of pasture in the rotation), including the traditional rice and pasture rotation phases (RP and RP2) and RCP, cluster in one group. A second group consists of rotations that include crops and short pasture (RS, RC, and RSP, with 50% of time with rice), and a third group consisting of continuous rice (CR, with 100% time with rice) in an isolated position (Fig. 5). These results showed that rice rather than soil use intensity or cover crop shaped the soil bacterial community after a complete cycle. Physicochemical changes that are occurring in the soil due to different uses are conditioning these microbial communities. Fungal communities are a little different and cluster mostly according to previous crop/pasture present in the rotation, although differences are poorly delineated after six years. Again, a greater definition in the identification of fungal taxa by sequencing other regions can help to better delineate these communities. The three groups clustered according to rice, pasture (one to three years long), or crops (soybean/sorghum) as antecessors (Fig. 6). This result is consistent with several studies that point out that plant type affect soil fungal community composition (Schmidt et al., 2019). Fungi depends more on plant products and their biotrophic interactions with different plant species because a large proportion of fungi interact narrowly as pathogens, saprotrophs or mutualists of plants (Millard and Singh, 2010; Schmidt et al., 2019). A more complete study, better delineating the taxa and making associations at the trophic or guild level of the fungi, is desirable to better characterize the communities and relationships existing in this group according to the rotation studied.

Some taxa were identified as biomarkers by LEfSe analysis for some of the rotations studied. While in the lax rotations (RP and RP2) *Proteobacteria* and *Actinobacteria* are the associated biomarkers, in intensive rotations (CR) there is a greater diversity of biomarkers, including *Actinobacteria*, *Proteobacteria* and *Chloroflexi* and Archaea. Archaeal communities including methanogenic *Methanosarcinaceae* are distinctive in irrigated rice field soils from most geographic regions (Fernández et al. 2013). Biomarkers found for RSP were mostly *Firmicutes* (*Ruminococcaceae*, *Paenibacillaceae* and *Ammoniphilus*), but reasons for this finding are unknown at this moment. Garaycochea et al. (2020) reported a higher abundance of *Firmicutes* in grassland soils with higher water holding capacity and permanent wilt point. LEfSe analysis for fungi failed to discriminate biomarkers and a more detailed study is needed to determine fungal indicator taxa for the rotations studied.

## Acknowledgements

I would like to thank Fernando Escalante for technical assistance in the project and the Rice Management Section personnel of INIA Treinta y Tres, for assistance provided during field sampling. This work was supported by the Instituto Nacional de Investigación Agropecuaria, Uruguay [Proyecto INIA AZ_40].

