## Supplemental Material for "Microbial community structure in rice, crops, and pastures rotation systems with different intensification levels in the temperate region of Uruguay"

Sebastián Martínez

**Table S1.** Relative abundance of the 20 most abundant bacterial taxa of classified sequences.

| Relative abundance | Taxa | Phylum |
|---|---|---|
| 4,90 | _Bacillus | Firmicutes |
| 3,21 | _Bacillus aryabhattai | Firmicutes |
| 2,76 | _uncultured Prosthecobacter sp. | Verrucomicrobia |
| 2,75 | _uncultured Conexibacteraceae bacterium | Actinobacteria |
| 2,64 | _uncultured Conexibacter sp. | Actinobacteria |
| 2,14 | _Nocardioides sp. | Actinobacteria |
| 2,13 | _Acidothermus | Actinobacteria |
| 1,50 | _Bradyrhizobium | Proteobacteria |
| 1,23 | _Bacillus | Firmicutes |
| 1,10 | _Pseudolabrys_uncultured bacterium | Proteobacteria |
| 1,03 | _Bacillus | Firmicutes |
| 1,02 | _Nocardioidaceae | Actinobacteria |
| 0,99 | _Candidatus Solibacter | Acidobacteria |
| 0,97 | _uncultured Sphingomonadaceae bacterium | Proteobacteria |
| 0,94 | _Streptomyces | Actinobacteria |
| 0,91 | _Terrabacter_uncultured bacterium | Actinobacteria |
| 0,81 | _Mycobacterium | Actinobacteria |
| 0,81 | _uncultured Rubrobacteria | Actinobacteria |
| 0,77 | _Xanthobacteraceae_uncultured forest soil bacterium | Proteobacteria |
| 0,76 | _Streptomyces | Actinobacteria |

**Table S2.** Relative abundance of the 20 most abundant fungal taxa of classified sequences.

| **Relative abundance.** | **Taxa** | **Orden** |
|---|---|---|
| 20,99 | _Fusarium oxysporum | Ascomycota |
| 11,97 | _Aspergillaceae | Ascomycota |
| 11,14 | _Chaetomium globosum | Ascomycota |
| 10,03 | _Fungi | |
| 5,40 | _Cucurbitariaceae; uncultured fungus | Ascomycota |
| 5,29 | _Talaromyces purpureogenus | Ascomycota |
| 3,87 | _Neophaeosphaeria; uncultured fungus | Ascomycota |
| 3,62 | _Fungi | |
| 2,62 | _Rhizopus oryzae | Mucoromycotina |
| 1,73 | _Alternaria alternata | Ascomycota |
| 1,63 | _Cladosporium | Ascomycota |
| 1,39 | _Saitozyma; uncultured Tremellaceae | Basidiomycota |
| 1,35 | _Plectosphaerellaceae | Ascomycota |
| 1,07 | _Arthopyreniaceae sp. GMG_P1 | Ascomycota |
| 1,00 | _Mortierella; uncultured fungus | Mortierellomycotina |
| 0,88 | _Fungi | |
| 0,86 | _Camarops | Ascomycota |
| 0,79 | _Pleosporales | Ascomycota |
| 0,77 | _Fungi | |
| 0,72 | _Athelia rolfsii | Basidiomycota |

**Table S3.** Pearsons's correlations and P-value of total Fungal classes, total Archaea phyla, and total Bacteria phyla with physicochemical parameters analyzed. Statistically significant correlations in bold ($P < 0.05$).

| Parameter | Group | Correlation | P value |
|---|---|---|---|
| | | **Fungi** | |
| N | Aphelidea | 0.38 | 0.091 |
| C | Aphelidea | 0.27 | 0.236 |
| CN | Aphelidea | -0.08 | 0.723 |
| pH | Aphelidea | -0.37 | 0.095 |
| P | Aphelidea | 0.24 | 0.304 |
| K | Aphelidea | 0.095 | 0.681 |
| Mg | Aphelidea | -0.11 | 0.643 |
| N | Ascomycota | 0.08 | 0.735 |
| C | Ascomycota | 0.11 | 0.636 |
| CN | Ascomycota | 0.19 | 0.412 |
| pH | Ascomycota | 0.15 | 0.513 |
| P | Ascomycota | 0.07 | 0.779 |
| K | Ascomycota | -0.47 | **0.032** |
| Mg | Ascomycota | -0.47 | **0.029** |
| N | Basidiomycota | 0.26 | 0.262 |
| C | Basidiomycota | 0.19 | 0.388 |
| CN | Basidiomycota | 0.01 | 0.951 |
| pH | Basidiomycota | -0.29 | 0.208 |
| P | Basidiomycota | 0.29 | 0.187 |
| K | Basidiomycota | 0.25 | 0.270 |
| Mg | Basidiomycota | 0.19 | 0.402 |
| N | Blastocladiomycota | -0.02 | 0.925 |
| C | Blastocladiomycota | 0.01 | 0.952 |
| CN | Blastocladiomycota | 0.18 | 0.434 |
| pH | Blastocladiomycota | -0.025 | 0.939 |
| P | Blastocladiomycota | -0.27 | 0.229 |
| K | Blastocladiomycota | -0.20 | 0.376 |
| Mg | Blastocladiomycota | -0.21 | 0.352 |
| N | Chytridiomycota | -0.01 | 0.954 |
| C | Chytridiomycota | -0.07 | 0.776 |
| CN | Chytridiomycota | -0.13 | 0.602 |
| pH | Chytridiomycota | -0.25 | 0.282 |
| P | Chytridiomycota | 0.43 | **0.050** |
| K | Chytridiomycota | -0.29 | 0.198 |
| Mg | Chytridiomycota | -0.39 | 0.079 |
| N | Olpidiomycota | -0.25 | 0.277 |
| C | Olpidiomycota | -0.26 | 0.262 |
| CN | Olpidiomycota | -0.189 | 0.446 |
| pH | Olpidiomycota | -0.05 | 0.835 |
| P | Olpidiomycota | -0.16 | 0.497 |
| K | Olpidiomycota | -0.10 | 0.666 |
| Mg | Olpidiomycota | -0.06 | 0.812 |
| N | Cryptomycota | 0.22 | 0.346 |
| C | Cryptomycota | 0.27 | 0.234 |
| CN | Cryptomycota | 0.40 | 0.071 |

| | | | |
|---|---|---|---|
| pH | Cryptomycota | -0.03 | 0.888 |
| P | Cryptomycota | -0.34 | 0.129 |
| K | Cryptomycota | 0.05 | 0.814 |
| Mg | Cryptomycota | -0.05 | 0.842 |
| N | Glomeromycota | -0.1752 | 0.447 |
| C | Glomeromycota | -0.2353 | 0.304 |
| CN | Glomeromycota | -0.2888 | 0.204 |
| pH | Glomeromycota | -0.4190 | 0.059 |
| P | Glomeromycota | 0.4650 | **0.034** |
| K | Glomeromycota | 0.3511 | 0.119 |
| Mg | Glomeromycota | 0.2214 | 0.335 |
| N | Incertae sedis | -0.1080 | 0.641 |
| C | Incertae sedis | -0.1134 | 0.625 |
| CN | Incertae sedis | -0.0429 | 0.853 |
| pH | Incertae sedis | 0.0122 | 0.958 |
| P | Incertae sedis | -0.2499 | 0.275 |
| K | Incertae sedis | -0.3320 | 0.141 |
| Mg | Incertae sedis | -0.0729 | 0.753 |
| N | Mucoromycota | -0.1761 | 0.445 |
| C | Mucoromycota | -0.1156 | 0.618 |
| CN | Mucoromycota | 0.1074 | 0.643 |
| pH | Mucoromycota | 0.2123 | 0.355 |
| P | Mucoromycota | -0.1544 | 0.504 |
| K | Mucoromycota | -0.1807 | 0.433 |
| Mg | Mucoromycota | -0.1398 | 0.546 |
| N | Mortierellomycota | 0.0957 | 0.679 |
| C | Mortierellomycota | 0.1064 | 0.646 |
| CN | Mortierellomycota | 0.1531 | 0.508 |
| pH | Mortierellomycota | -0.2037 | 0.376 |
| P | Mortierellomycota | -0.0938 | 0.686 |
| K | Mortierellomycota | 0.3283 | 0.146 |
| Mg | Mortierellomycota | -0.0617 | 0.790 |
| N | Neocallimastigomycota | -0.0403 | 0.862 |
| C | Neocallimastigomycota | -0.1247 | 0.590 |
| CN | Neocallimastigomycota | -0.2706 | 0.236 |
| pH | Neocallimastigomycota | -0.3924 | 0.079 |
| P | Neocallimastigomycota | 0.1251 | 0.589 |
| K | Neocallimastigomycota | 0.0536 | 0.818 |
| Mg | Neocallimastigomycota | -0.0515 | 0.825 |
| N | Nuclearida | -0.0333 | 0.886 |
| C | Nuclearida | -0.0617 | 0.790 |
| CN | Nuclearida | -0.1240 | 0.592 |
| pH | Nuclearida | -0.0629 | 0.786 |
| P | Nuclearida | 0.4564 | **0.038** |
| K | Nuclearida | 0.0138 | 0.953 |
| Mg | Nuclearida | -0.1899 | 0.409 |
| N | Unidentified | -0.1942 | 0.399 |
| C | Unidentified | -0.1342 | 0.562 |
| CN | Unidentified | -0.0075 | 0.974 |
| pH | Unidentified | 0.5183 | **0.016** |
| P | Unidentified | -0.0061 | 0.979 |
| K | Unidentified | -0.4039 | 0.069 |
| Mg | Unidentified | 0.1644 | 0.476 |

| | | | |
|---|---|---|---|
| N | Zoophagomycota | -0.0783 | 0.736 |
| C | Zoophagomycota | -0.0967 | 0.677 |
| CN | Zoophagomycota | -0.0738 | 0.751 |
| pH | Zoophagomycota | -0.2949 | 0.195 |
| P | Zoophagomycota | 0.4267 | **0.054** |
| K | Zoophagomycota | 0.1633 | 0.479 |
| Mg | Zoophagomycota | 0.2258 | 0.325 |
| **Archaea/Bacteria** | | | |
| N | Actinobacteria | -0.1234 | 0.594 |
| C | Actinobacteria | -0.1065 | 0.646 |
| CN | Actinobacteria | -0.0078 | 0.973 |
| pH | Actinobacteria | 0.2279 | 0.320 |
| P | Actinobacteria | -0.4270 | 0.054 |
| K | Actinobacteria | -0.4987 | **0.021** |
| Mg | Actinobacteria | 0.0026 | 0.991 |
| N | Firmicutes | 0.1169 | 0.614 |
| C | Firmicutes | 0.0740 | 0.749 |
| CN | Firmicutes | 0.1429 | 0.537 |
| pH | Firmicutes | 0.1319 | 0.569 |
| P | Firmicutes | 0.0227 | 0.922 |
| K | Firmicutes | 0.0473 | 0.839 |
| Mg | Firmicutes | -0.2963 | 0.192 |
| N | Proteobacteria | -0.3662 | 0.103 |
| C | Proteobacteria | -0.2623 | 0.251 |
| CN | Proteobacteria | -0.0403 | 0.862 |
| pH | Proteobacteria | 0.3326 | 0.141 |
| P | Proteobacteria | -0.5746 | **0.006** |
| K | Proteobacteria | -0.3831 | 0.087 |
| Mg | Proteobacteria | 0.0151 | 0.948 |
| N | Chloroflexi | -0.1922 | 0.404 |
| C | Chloroflexi | -0.2558 | 0.263 |
| CN | Chloroflexi | -0.3844 | 0.085 |
| pH | Chloroflexi | -0.1968 | 0.392 |
| P | Chloroflexi | -0.0897 | 0.699 |
| K | Chloroflexi | -0.1958 | 0.395 |
| Mg | Chloroflexi | 0.1310 | 0.571 |
| N | Acidobacteria | -0.1987 | 0.388 |
| C | Acidobacteria | -0.1727 | 0.454 |
| CN | Acidobacteria | -0.1636 | 0.478 |
| pH | Acidobacteria | -0.1137 | 0.624 |
| P | Acidobacteria | 0.0591 | 0.799 |
| K | Acidobacteria | 0.1183 | 0.609 |
| Mg | Acidobacteria | 0.1501 | 0.516 |
| N | Verrucomicrobia | -0.1871 | 0.417 |
| C | Verrucomicrobia | -0.1643 | 0.477 |
| CN | Verrucomicrobia | -0.2124 | 0.355 |
| pH | Verrucomicrobia | -0.1605 | 0.487 |
| P | Verrucomicrobia | 0.4532 | **0.039** |
| K | Verrucomicrobia | 0.3885 | 0.082 |
| Mg | Verrucomicrobia | 0.1403 | 0.442 |
| N | Euryarchaeota | -0.1345 | 0.561 |
| C | Euryarchaeota | -0.1208 | 0.602 |
| CN | Euryarchaeota | -0.1812 | 0.432 |

| | | | |
|---|---|---|---|
| pH | Euryarchaeota | -0.0858 | 0.712 |
| P | Euryarchaeota | -0.2239 | 0.329 |
| K | Euryarchaeota | -0.0335 | 0.885 |
| Mg | Euryarchaeota | 0.1373 | 0.553 |
| N | Planctomycetes | 0.2455 | 0.284 |
| C | Planctomycetes | 0.2247 | 0.328 |
| CN | Planctomycetes | 0.1870 | 0.417 |
| pH | Planctomycetes | -0.1578 | 0.494 |
| P | Planctomycetes | 0.1443 | 0.533 |
| K | Planctomycetes | 0.4008 | 0.072 |
| Mg | Planctomycetes | -0.0704 | 0.762 |
| N | Bacteroidetes | -0.1584 | 0.493 |
| C | Bacteroidetes | -0.1558 | 0.499 |
| CN | Bacteroidetes | -0.1506 | 0.515 |
| pH | Bacteroidetes | 0.3124 | 0.168 |
| P | Bacteroidetes | 0.1930 | 0.402 |
| K | Bacteroidetes | -0.1656 | 0.473 |
| Mg | Bacteroidetes | -0.1903 | 0.409 |
| N | Synergistetes | -0.4885 | **0.025** |
| C | Synergistetes | -0.4625 | **0.035** |
| CN | Synergistetes | -0.2183 | 0.342 |
| pH | Synergistetes | 0.1799 | 0.435 |
| P | Synergistetes | - 0.228 | 0.318 |
| K | Synergistetes | -0.3749 | 0.094 |
| Mg | Synergistetes | -0.0336 | 0.885 |
| N | Armatimonadetes | -0.22 | 0.336 |
| C | Armatimonadetes | -0.23 | 0.306 |
| CN | Armatimonadetes | -0.24 | 0.303 |
| pH | Armatimonadetes | -0.12 | 0.592 |
| P | Armatimonadetes | 0.15 | 0.521 |
| K | Armatimonadetes | -0.19 | 0.393 |
| Mg | Armatimonadetes | -0.16 | 0.493 |
| N | Spirochaetes | -0.38 | 0.091 |
| C | Spirochaetes | -0.38 | 0.085 |
| CN | Spirochaetes | -0.33 | 0.150 |
| pH | Spirochaetes | 0.35 | 0.122 |
| P | Spirochaetes | 0.24 | 0.294 |
| K | Spirochaetes | 0.09 | 0.676 |
| Mg | Spirochaetes | -0.11 | 0.635 |
| N | WPS | 0.03 | 0.889 |
| C | WPS-2 | 0.01 | 0.949 |
| CN | WPS-2 | -0.03 | 0.913 |
| pH | WPS-2 | -0.31 | 0.178 |
| P | WPS-2 | 0.02 | 0.941 |
| K | WPS-2 | -0.26 | 0.246 |
| Mg | WPS-2 | -0.17 | 0.472 |
| N | Thermotogae | -0.05 | 0.845 |
| C | Thermotogae | -0.06 | 0.808 |
| CN | Thermotogae | -0.06 | 0.782 |
| pH | Thermotogae | 0.13 | 0.582 |
| P | Thermotogae | 0.17 | 0.454 |
| K | Thermotogae | -0.06 | 0.811 |
| Mg | Thermotogae | -0.33 | 0.139 |

| | | | |
|---|---|---|---|
| N | Gemmatimonadetes | -0.08 | 0.733 |
| C | Gemmatimonadetes | -0.08 | 0.733 |
| CN | Gemmatimonadetes | -0.17 | 0.461 |
| pH | Gemmatimonadetes | 0.18 | 0.442 |
| P | Gemmatimonadetes | -0.43 | 0.054 |
| K | Gemmatimonadetes | -0.09 | 0.669 |
| Mg | Gemmatimonadetes | 0.03 | 0.894 |
| N | Crenarchaeota | -0.22 | 0.343 |
| C | Crenarchaeota | -0.19 | 0.397 |
| CN | Crenarchaeota | -0.24 | 0.286 |
| pH | Crenarchaeota | -0.05 | 0.819 |
| P | Crenarchaeota | -0.22 | 0.342 |
| K | Crenarchaeota | 0.02 | 0.934 |
| Mg | Crenarchaeota | 0.26 | 0.262 |
| N | Thaumarchaeota | 0.31 | 0.166 |
| C | Thaumarchaeota | 0.22 | 0.346 |
| CN | Thaumarchaeota | 0.08 | 0.735 |
| pH | Thaumarchaeota | -0.46 | **0.038** |
| P | Thaumarchaeota | 0.37 | 0.094 |
| K | Thaumarchaeota | 0.24 | 0.296 |
| Mg | Thaumarchaeota | 0.07 | 0.749 |
| N | Cyanobacteria | -0.02 | 0.922 |
| C | Cyanobacteria | -0.01 | 0.959 |
| CN | Cyanobacteria | 0.10 | 0.674 |
| pH | Cyanobacteria | 0.13 | 0.571 |
| P | Cyanobacteria | -0.36 | 0.113 |
| K | Cyanobacteria | -0.46 | **0.035** |
| Mg | Cyanobacteria | -0.35 | 0.123 |
| N | Nitrospirae | -0.26 | 0.256 |
| C | Nitrospirae | -0.32 | 0.162 |
| CN | Nitrospirae | -0.47 | **0.031** |
| pH | Nitrospirae | 0.14 | 0.547 |
| P | Nitrospirae | -0.09 | 0.702 |
| K | Nitrospirae | -0.12 | 0.597 |
| Mg | Nitrospirae | 0.19 | 0.417 |

**Fig S1.** Crop rotation systems, annual and seasonal schedules at the Long-term experiment of Unidad Experimental de Paso de la Laguna, Treinta y Tres, Uruguay.

| | | | Season | | | | | | | | | | | |
|---|---|---|---|---|---|---|---|---|---|---|---|---|---|---|
| Rotation* | | | 1 | | 2 | | 3 | | 4 | | 5 | | 6 | |
| **Nº** | **Name** | **Code** | **SS** | **AW** | **SS** | **AW** | **SS** | **AW** | **SS** | **AW** | **SS** | **AW** | **SS** | **AW** |
| 1 | **Continuous rice** | **CR** | Rice | CC Ta | | | | | | | | | | |
| 2 | **Rice and crops** | **RC** | Rice | CC Ec | Soybean | CC Ec | Rice | CC Ec | Sorghum | CC Ec | | | | |
| 3 | **Rice and short pasture** | **RSP** | Rice | P Rc-Ry | P | P | | | | | | | | |
| 4 | **Rice and long pasture** | **RP** | Rice | CC Ry | Rice | P wc-L-F | P | P | P | P | P | P | | |
| 5 | **Rice, crop and pasture** | **RCP** | Rice | CC Ry | Soybean | CC Ry | Soybean | CC Ta | Rice | P L-Ry | P | P | P | P |
| 6 | **Rice and soybean** | **RSP** | Rice | CC Ry | Soybean | CC Ta | | | | | | | | |

* SS= spring-summer, AW= autumn-winter, CC=cover crop, Ry=ryegrass (*Lolium multiflorum*), Rc=red clover (*Trifolium pratense*), wc=white clover, L=*Lotus* sp., P=pasture (*Festuca arundinacea*, *Trifolium repens* and *Lotus corniculatus*).

**Fig S2.** Rarefaction curves of 16S rDNA gene showing archaeal/bacterial OTUs.

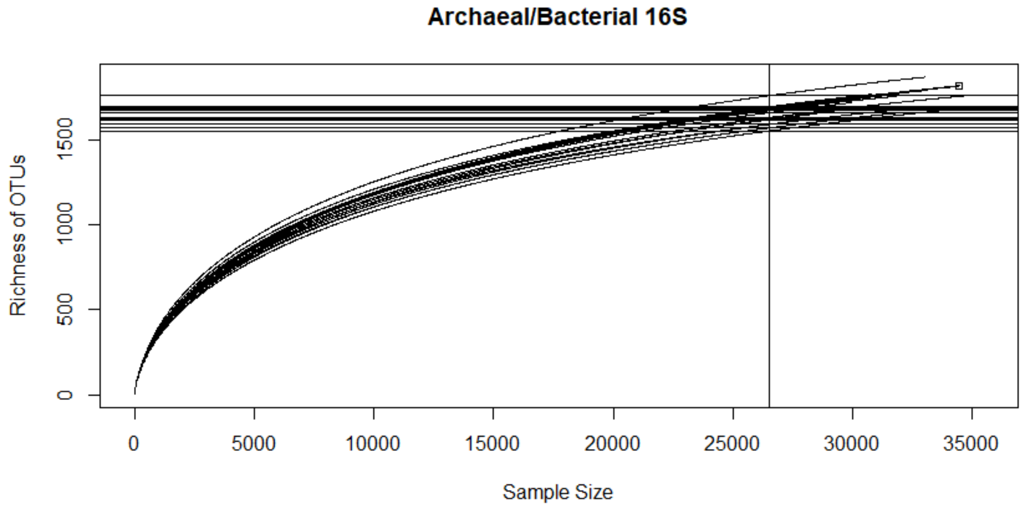

**Fig S3.** Rarefaction curves of 18S rDNA gene showing fungal OTUs.

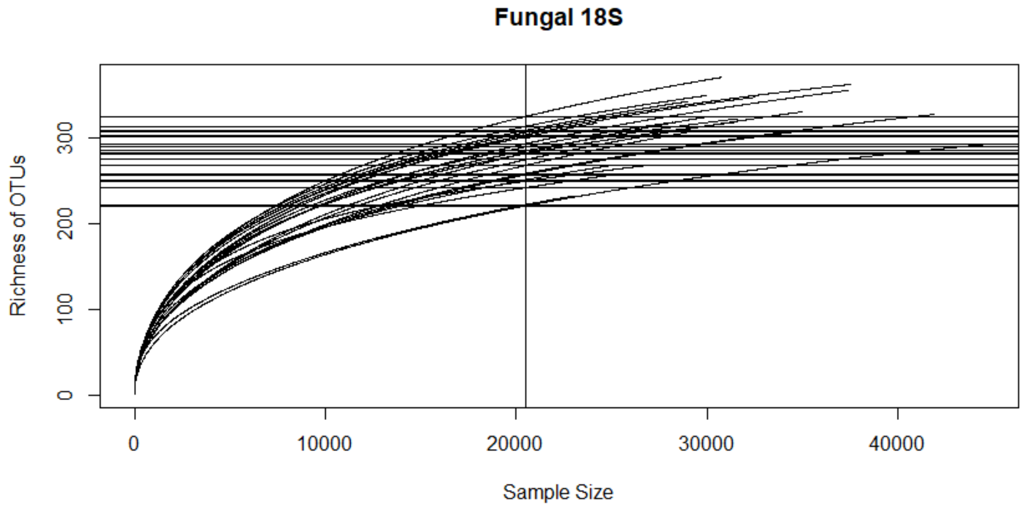

**Fig S4.** Total number of the 97% sequence similarity OTUs for fungal (18S) and bacterial/archaeal (16S) taxa. Assignment to fungal classes or bacterial phyla is according to UNITE database based on closely related sequences found.

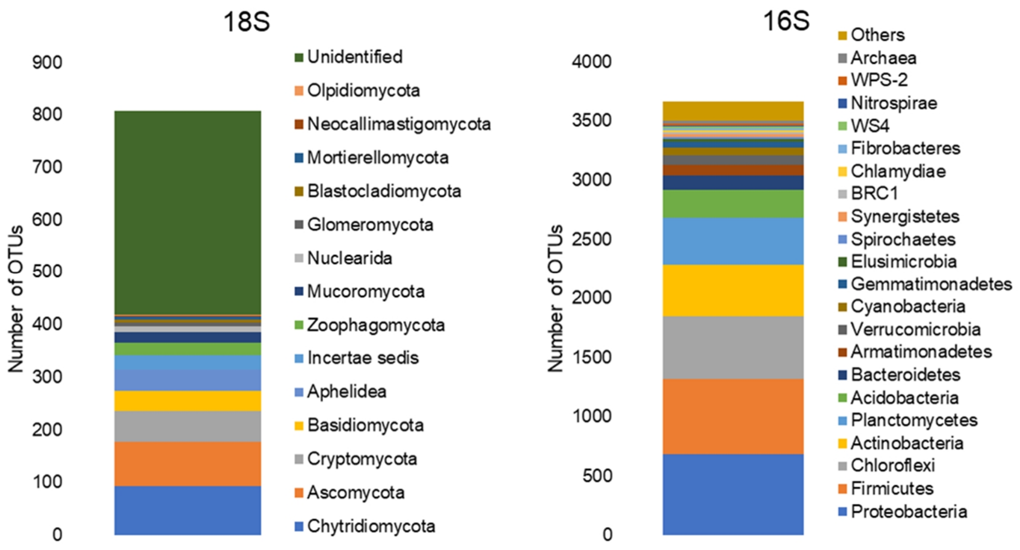

**Fig S5.** Relative abundance found of archaeal phyla *Crenarchaeota* and bacterial phyla *Cyanobacteria* and *Nitrospira* according with rotation studied. Rotations: CR=continuous rice, RC=rice and crops, RSP=rice and short pasture, RP=rice and pasture (after pasture), RP2=rice and pasture (after rice), RCP=rice, crops and pasture, and RS=rice and soybean (see Fig. 1). Bars represents standard errors based in three replications.

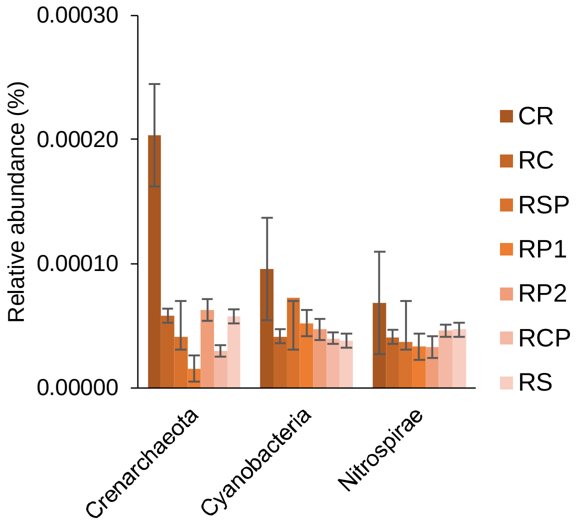

**Fig. S6.** LEfSe results based on $P < 0.05$ and an LDA score > 2.5 revealed bacteria biomarkers from phylum level to genus level that were sensitive to the different rotations studied. Only the bacterial lineages that showed any significant responses to rotation are shown. Rotations: CR=continuous rice, RC=rice and crops, RSP=rice and short pasture, RP=rice and pasture (after pasture), RP2=rice and pasture (after rice), RCP=rice, crops and pasture, and RS=rice and soybean

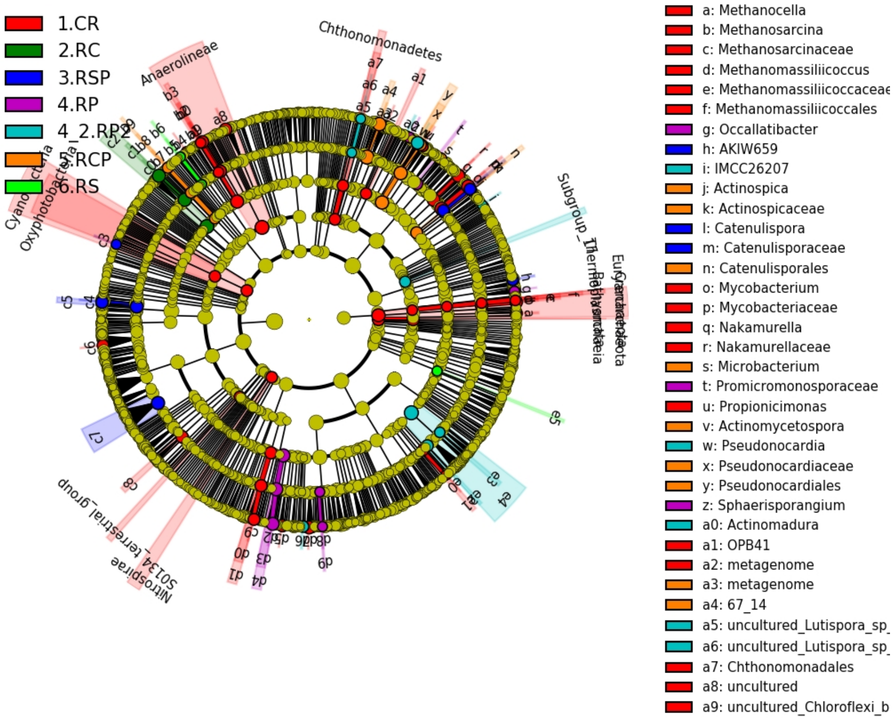

**Fig S7.** Linear discriminant analysis Effect Size (LEfSe) (log10 LDA score) of fungal OTUs, which most likely explain differences between rice rotations after a complete cycle of six years. Fungal OTUs are classified at the highest resolvable taxonomic level. Rotations: CR= continuous rice, RP= rice and pasture (after pasture), RP2= rice and pasture (after rice), and RS= rice and soybean.

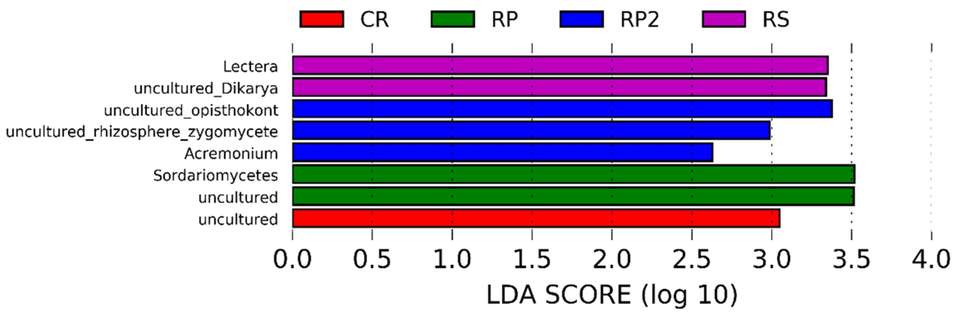
